# Integrative morphology and phylogenetics of Arcellidae (Amoebozoa:Arcellinida), with redescription of *Arcella leidyana* and *Arcella artocrea* and description of *Galeripora purdoni* sp. nov

**DOI:** 10.64898/2026.08.19.745684

**Authors:** Bruce D. S. Taylor, Alfredo L. Porfirio-Sousa, Robert E. Jones, Carl Seaquist, Ferry J. Siemensma, Elias Taylor, Alexander K. Tice

## Abstract

Arcellidae is a family of testate amoebae within Arcellinida (Amoebozoa), comprising three recognized genera: *Arcella*, *Galeripora*, and *Antarcella*. Although species in the family have been studied for nearly two centuries, many historically described taxa and major morphological groups remain unsampled at the molecular level. Here, we provide a comprehensive review of Arcellidae and generate new cytochrome c oxidase subunit I (COI) sequences for arcellid species from Canadian peatlands, focusing on tall-shelled *Arcella* historically classified in section Altae sensu Deflandre. COI phylogenetic analyses recover a strongly supported monophyletic clade corresponding to North American representatives of Altae, providing the first molecular corroboration of this morphologically defined group. Within this clade, we redescribe *Arcella leidyana* based on modern material from Eeyou Istchee (Quebec). We further describe *Galeripora purdoni* sp. nov. from a calcareous fen in eastern Ontario, representing a novel terrestrial lineage within the genus, and redescribe *Galeripora artocrea*, which we transfer to *Arcella* based on congruent molecular and morphological evidence. Phylogenomic analyses of Arcellidae isolates from the Protist 10,000 Genomes Project reveal an additional deep lineage basal to *Arcella* and *Galeripora*. Together, these results highlight hidden diversity and demonstrate the importance of integrative approaches for resolving arcellid systematics and refining its classification.

## 1. Introduction

Arcellidae Ehrenberg, 1830 is a family of shelled (testate) amoebae in the order Arcellinida (Amoebozoa), comprising species whose shells (tests) are composed of secreted organic material arranged in hexagonal areoles, also called building units (Meisterfeld, 2002; Tsyganov et al., 2016). Currently, three genera are recognized within the family: *Arcella* Ehrenberg, 1830, *Galeripora* González-Miguéns et al., 2022 and *Antarcella* Deflandre, 1928, emend. Deflandre, 1953. While the monophyly of the family has been confirmed (González-Miguéns et al., 2022; Useros et al., 2023), relationships among its constituent lineages remain unsettled, and many historically described taxa have yet to be evaluated with molecular data.

Species delimitation of Arcellidae has traditionally relied on shell morphology, such as general shape, height, aperture shape, surface ornamentation and number of nuclei (Meisterfeld, 2002). Based on its morphology, Deflandre (1928) divided the genus *Arcella* into four sections. Molecular phylogenetics has since shown that three of these sections—Vulgares, Carinatae, and Aplanatae—are non-monophyletic, underscoring the extent of morphological convergence and homoplasy in the group (González-Miguéns et al., 2022). The fourth section, Altae, comprising species with relatively tall shells and steeply angled sides, has not yet been evaluated with molecular evidence.

Here, we provide a comprehensive review of the literature on Arcellidae, together with an integrative phylogenetic and morphological investigation based on new cytochrome c oxidase subunit I (COI) sequences, expanded phylogenomic sampling, and SEM-based shell analyses. A particular focus is on North American members of Deflandre’s (1928) section Altae, a morphologically distinctive group of tall-shelled species possessing crenulated apertures, originally treated by Joseph Leidy as morphotypes of *Arcella mitrata*. This group has not previously been evaluated with molecular evidence.

Our COI analyses demonstrate that North American representatives of Altae form a well-supported monophyletic clade within *Arcella*. Within this clade, we redescribe *Arcella leidyana* based on modern material from Eeyou Istchee (Quebec), Canada. Outside this group, our results support returning *Galeripora artocrea* to *Arcella*, and we describe *Galeripora purdoni* sp. nov. from a calcareous fen in eastern Ontario, Canada. Phylogenomic reconstruction further reveals a previously unrecognized deep lineage basal to both *Arcella* and *Galeripora*, refining the evolutionary framework of the family.

## 2. Materials and Methods

### 2.1 Study sites and sample collection

Samples for this study were collected between 2024 and 2025, at three locations in Canada (Table 1). At Mer Bleue, samples were gathered from two sites on a small dystrophic lake bounded on its southern edge by an extensive *Sphagnum* dome. The Eeyou Istchee sites lie within a vast subarctic coastal plain along the northeastern shore of James Bay, characterized by extensive wetlands, muskeg bogs, and sparse open taiga dominated by black spruce (*Picea mariana*). Samples were taken in July, 2025 from saturated scorpion moss (*Scorpidium* sp.) and *Sphagnum*. The site in Purdon Conservation Area is located within a protected wetland in Lanark, Ontario, a small calcareous fen fed by groundwater rich in dissolved minerals derived from limestone and dolostone bedrock. Samples were taken in September, 2025, alongside a public boardwalk, from leaf litter and woody debris strewn among hummocks of minerotrophic *Sphagnum*.

**Table 1.**
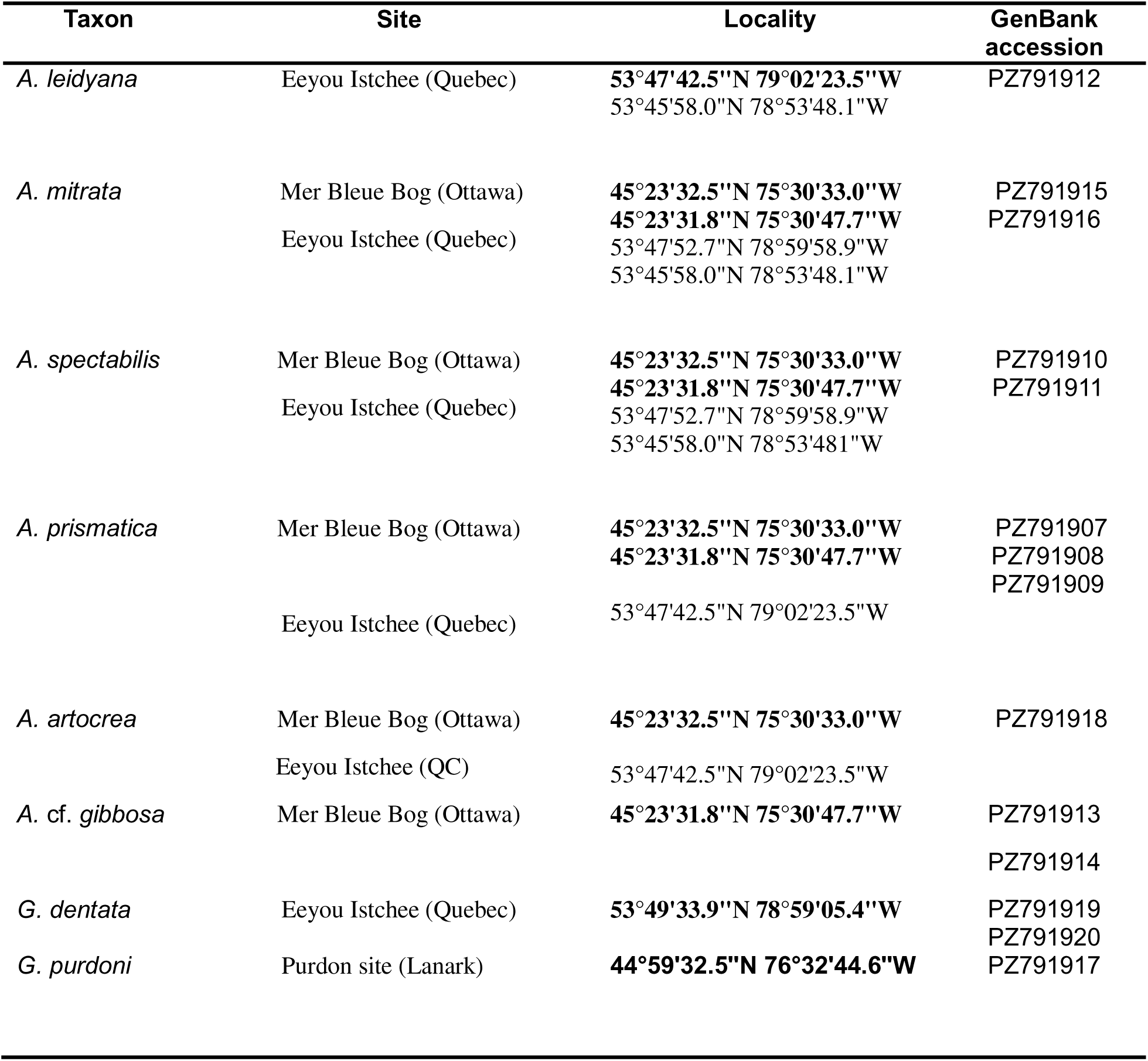
Localities of the investigated Arcellidae species and GenBank accession numbers for newly generated cytochrome c oxidase subunit I (COI) sequences. Coordinates shown in bold indicate the populations from which the COI sequences were generated.

**Table 2.** Morphometrics of the Arcellidae considered in the present study as shown in Figure 1. MD – maximum diameter; VD – ventral diameter; MH – maximum height; AD – aperture diameter; AH – aperture height; AD/VD - ratio of AD and VD; MD/MH – ratio of MD and MH; VD/MD – ratio of VD/MD; AH/MD – ratio of AH/MD; N of Ribs – number of ribs; AoS – angle of sides. Parentheses indicate the number of individuals measured (n).

| Taxon | MD | VD | MH | AD | AH | AD/VD | MD/MH | VD/M |
| --- | --- | --- | --- | --- | --- | --- | --- | --- |
| *A. leidyana* | 116–152(24) | 120–150(47) | 102–134(19) | 40–67(34) | 9–19(13) | 0.38–0.47(33) | 1.1–1.3(19) | 0.9–1.1 |
| *A. mitrata* | 115–150(31) | 76–113(30) | 112–144(29) | 32–43(6) | 25–38(6) | 0.33–0.40(3) | 0.9–1.1(27) | 0.5–0.82( |
| *A. spectabilis*<br>(large) | 110–145(45) | 87–117(39) | 93–139(33) | 26–33(12) | 17–34(25) | 0.20–0.25(9) | 1.0–1.22(30) | 0.7–0.9 |
| *A. spectabilis*<br>(small) | 80–100(79) | 59–80(64) | 75–102(40) | 23–30(24) | - | 0.31–0.45(24) | 0.91–1.15(40) | 0.67–0.8 |
| *A. prismatica* | 55–97(37) | 81–115(87) | 61–87(34) | 27–33(41) | 9–13(16) | 0.27–0.38(34) | 1–1.5(34) | 1–1.8 |
| *A. artocrea*<br>(*Mer Bleue*) | 130–159(84) | - | 41–56(15) | 19–33(66) | - | 0.13–0.23(66) | 0.29–0.36(15) | - |
| *A. artocrea*<br>(Chisasibi) | 141–178(48) | - | - | 22–31(28) | - | 0.12–0.2(28) | - | - |
| *G. purdoni* | 178–253(56) | - | 77–101(7) | 36–56(47) | 34–43(7) | 0.18–0.24(47) | 2.1–2.8(7) | - |

Sampling procedures were the same at all sites, except Purdon Fen. For each sample, approximately 100 mL of surface water and associated vegetation were collected in a glass jar. Multiple samples were then screened for the presence of Arcellidae. Aliquots of liquid (1-3 mL) were extracted from each sample jar, transferred to a 70 mm Petri dish and examined at 40× under a stereo microscope. Arcellid shells were isolated with a glass micropipette for further analysis. Some specimens were deposited in 1.5 mL micro-centrifuge tubes and set aside for morphometrics and SEM imaging. Others were washed in successive drops of distilled water, then placed in guanidine thiocyanate (TG) buffer solution and stored at -18°C for subsequent PCR.

The dry samples from Purdon Fen were placed in 90 mm Petri dishes, saturated with water, and allowed to soak overnight. Suspensions of water and loose particulate matter were then extracted with a 3 mL transfer pipette and deposited into 70 mm Petri dishes. These were further diluted and examined at 40× magnification under a stereo microscope. Specimens of *Galeripora* were picked out with small-diameter pipettes and transferred to concave (cavity) slides for measurement and observation by light microscopy. The contents of the slides were then washed into fresh Petri dishes. Shells containing live amoebae were placed in vials of TG solution and set aside for subsequent PCR. Empty and encysted shells were prepared for SEM imaging as described below.

### 2.2 Morphometrics and Imaging

Shell measurements were carried out using Motic 310E compound microscope, equipped with an AmScope MU300 camera, and ToupView 4.1 software, 400× magnification. Key shell dimensions followed the scheme in Figure 1. After measurement, shells were washed into Petri dishes, and representative shells were isolated with a glass micropipette, mounted on two-sided adhesive disks affixed to aluminum SEM stubs and positioned with a single-hair eyelash manipulator. Specimens were air-dried under cover and then sputter-coated with gold-palladium for 25 seconds in a Denton Desk II sputter coater. Imaging was performed with a Thermo Scientific Apreo II field emission SEM at the Canadian Museum of Nature. SEM images were acquired with both secondary electron (ETD) and backscattered electron (BSE) detectors at various magnifications, accelerating voltages, and beam currents; additional details are provided in the figure captions.

**Figure 1.**
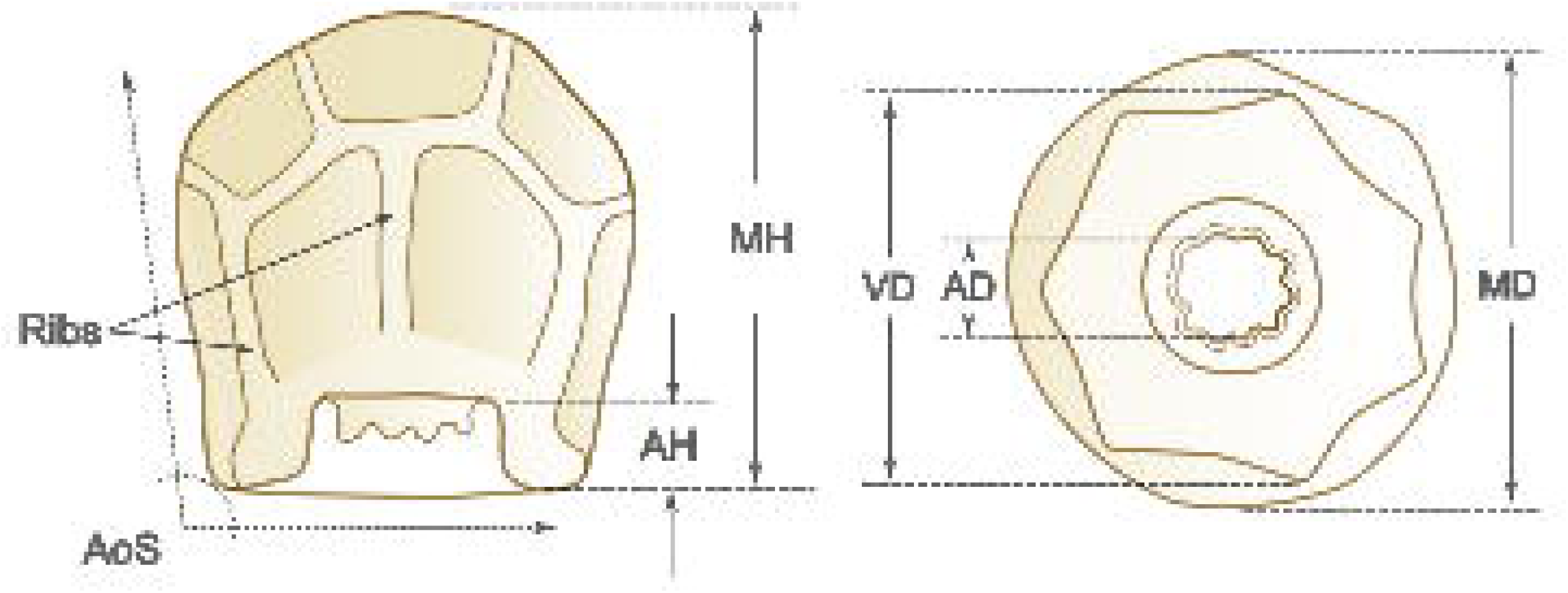
Principal dimensions used for morphometrics. AD = aperture diameter; AH = aperture height; AoS = angle of Sides; MD = maximum diameter; MH = maximum height; VD = ventral diameter (diameter at the ventral face).

### 2.2 DNA extraction, PCR and sequencing

Cells collected in the field were deposited individually in cryovials containing 100 μL aliquots of guanidine thiocyanate (TG) solution, prepared according to Duckert et al. (2018). These were kept at -18°C until PCR amplification and sequencing were performed. For PCR and sequencing, two different protocols were followed.

A first series of partial COI gene sequences was obtained in February, 2025 for *Arcella prismatica*, *A. leidyana* and *A. spectabilis* (large morphotype). PCR amplification and Sanger sequencing were performed at the University of Guelph Advanced Analysis Center (Genomics Facility). Amplification of the COI gene region was performed using the semi-nested protocol described in González-Miguéns et al. (2022). An initial round of amplification was done using the mitochondrial cytochrome c oxidase subunit I universal primer. This was followed by a second amplification, using *Arcella*-specific primers designed by González-Miguéns et al. (2022). Amplicons were sequenced by Sanger sequencing and results were checked against COI sequences of Arcellidae in GenBank to ensure that they clustered in the same family.

A second series of sequences was retrieved in September and October, 2025 for the species *Arcella prismatica*, *A. mitrata*, *A.* cf*. gibbosa*, *A. spectabilis* (small morphotype), *Galeripora dentata*, *G. artocrea*, and *G. purdoni* sp. nov. For amplification and sequencing of these individuals, a different procedure was followed. As with the previous group, specimens were placed in cryovials of TG solution and transferred to the University of Guelph Advanced Analysis Center (Genomics Facility) for PCR and sequencing. 14.25 µL of a multi-individual extraction from each species was used as input for a PCR protocol modified from (Useros et al., 2023). PCR was performed using a KAPA HiFi HotStart PCR Kit according to manufacturer instructions (KR0369 – v10.17) in two steps: 40 cycles with universal eukaryotic COI primers, LCO 1490 (GGTCAACAAATCATAAAGATATTGG) and HCO 2198 (TAAACTTCAGGGTGACCAAAAAATCA), followed by a 1:20 dilution, and then a further 45 cycles with the same primer set, for a total of 85 cycles. The PCR cycling conditions were unchanged from Useros et al.(2023): an initial denaturation at 96◦C for 5 min, followed by 40 cycles at 94◦C for 15 s, 40◦C for 15 s and 72◦C for 90 s and a final extension step at 72◦C for 10 min. An Agilent TapeStation 4150 was used to confirm the presence of products approximately 700 bp in size.

PCR products were cleaned using NucleoMag NGS Clean-up and Size Select® magnetic beads at a 0.8 dilution ratio. The cleaned PCR products were used as input for Oxford Nanopore long read sequencing on an ONT PromethION 24 with the Native Barcoding Kit 96 V14 (SQK-NBD114.96). Raw reads were filtered for 500-900 bp size, and the ONT wf-amplicon workflow (v1.1.4-g1cf4682) was used to generate consensus sequences and variants. Minimap2 (2.30-r1287) was used to align raw reads to the consensus sequences, and the correct consensus sequence was manually verified for each variant, discarding non-Arcellinida contaminant reads.

### 2.3 COI Phylogenetic Analyses

We compiled a cytochrome c oxidase subunit I (COI) dataset representing a broad sampling of Arcellidae family. These datasets were derived from NCBI, Protist 10,000 Genomes (P10K database, Gao et al., 2024), and previously constructed and curated datasets, aiming to construct a comprehensive Arcellidae datasets (González-Miguéns et al., 2022; Porfirio-Sousa et al., 2026; Useros et al., 2023). The raw COI dataset is presented in the Supporting Information (TableS1). The phylogenetic reconstructions were based on multiple sequence alignments (MSAs) generated using MAFFT v7.490 following command: mafft --auto --adjustdirectionaccurately input.fasta > output_aligned.fasta. Automated alignment trimming was performed with trimAl v1.5.rev1, using the command trimal -in input_aligned.fasta -out output_aligned_trimmed.fasta -keepheader -gt [threshold], where the gap threshold was set to 0.3. The COI-based phylogenetic tree was inferred from the trimmed alignments using the maximum likelihood method implemented in IQ-TREE v3.0.1, with ModelFinder for model selection and node support assessed via 1,000 SH-aLRT tests and 1,000 ultrafast bootstrap replicates. Support values of SH-aLRT/UFBoot ≥ 80/95 are considered indicative of strong support, following the recommendations of the original papers describing the development of these node support measures (Cornet and Baurain, 2022; Guindon et al., 2010). The analysis was executed with the command iqtree3 -s aligned_trimmed.fasta -alrt 1000 -bb 1000 -m MFP.

### 2.4 Phylogenomic Dataset Construction

We constructed our Arcellidae phylogenomic dataset from the genomic level assemblies available in the P10K database and that were previously assigned to Arcellidae family based on COI and SSU data (Porfirio-Sousa et al., 2026). Specifically, we used the database and tools provided by PhyloFisher v1.2.11 (Tice et al., 2021) following the detailed workflow available at https://thebrownlab.github.io/phylofisher-pages/detailed-example-workflow and the PhyloFisher protocol (Jones et al., 2024). Using the phylogenetically aware mode of fisher.py, we searched putative homologs of 240 target genes from the proteomes of 24 Arcellidae genomic level assemblies available in the P10K database (Gao et al., 2024). As queries, we used orthologs previously identified in the PhyloFisher database from the tubulinid amoebozoans *Arcella uspiensis* (intermedia), *Cryptodifflugia operculata*, and *Copromyxa protea*, which were employed in HMMER and BLAST searches executed by fisher.py (Camacho et al., 2009; Mistry et al., 2013). For each of the 240 putative orthologs, sequences recovered by BLAST were incorporated into their respective alignments using working_dataset_constructor.py. These alignments already included orthologs and paralogs from diverse eukaryotic taxa included in the database provided by PhyloFisher v1.2.11 (Tice et al., 2021) and a comprehensive sampling of Arcellinida taxa from previous phylogenomic studies (Kang et al., 2017; Lahr et al., 2019; Porfírio-Sousa et al., 2024). Homolog trees were inferred from the extended alignments using sgt_constructor.py, and each tree was manually inspected in ParaSorter to confirm correct ortholog and paralog assignments and to exclude any potential contaminant sequences from non-target eukaryotes. Final ortholog/paralog designations were applied to the PhyloFisher database using apply_to_db.py. To mitigate the effects of missing data in downstream analyses, we retained only orthologs present in at least 33% of the final taxon set. The resulting dataset was assembled using prep_final_dataset.py and matrix_constructor.py with default parameters. The final concatenated matrix used for phylogenetic analyses included 206 genes (57,156 amino acid sites) and was composed of 26 Arcellidae taxa alongside two *Netzelia* species included as outgroup (Supporting Information - Tables S2-S3).

### 2.5 Phylogenomic Analyses

We performed maximum likelihood (ML) phylogenetic reconstruction using our final matrix with IQ-TREE v3.0.1 (Wong et al., 2026). We first inferred a tree under the ELM+C20+F+G site heterogeneous model of evolution. Using this tree as a guide, we inferred another tree using the ELM+C60+F+G model of evolution in IQ-TREE3, collecting posterior mean site frequencies (PMSF) inferred from the dataset. We assessed topological support for the resulting tree with 100 non-parametric real bootstrap replicates under the ELM+C60+F+G+PMSF model of evolution in IQ-TREE3.

To assess phylogenetic relationships using combined evidence, the COI and PhyloFisher phylogenomic datasets were concatenated into a single supermatrix. The concatenated dataset comprised 58,644 aligned sites, including 1,488 COI nucleotide sites and 57,156 amino acid sites derived from protein coding genes found in the PhyloFisher database. Maximum-likelihood phylogenetic inference was conducted in IQ-TREE v3.1.1, applying separate partition-specific models (GTR+F+I+G4 for COI and ELM+C20+F+G for the PhyloFisher database partition). Node support was assessed using 1,000 ultrafast bootstrap replicates and 1,000 SH-aLRT replicates.

## 3. Results and Discussion

### 3.1 Review of the Arcellidae

#### 3.1.1 Early taxonomy

In 1877, Schulze erected the Arcellidae as a family of lobose *Rhizopoden* (testate amoebae with bluntly rounded pseudopods), comprising species whose tests were made entirely of “chitin” and also exhibited a visible “lattice structure” (*Gitterleistenwerk*) (Schulze, 1877). This “lattice structure” of the shell is now understood to be a mesh of discrete, hollow building units, produced as vesicles within the cell and secreted across the plasma membrane during ontogenesis (Netzel, 1975; Porfírio-Sousa A.L. and Lahr D.J.G., 2020).

The characterization of Arcellidae by shell composition remained in use throughout the 20^th^ century, with varying circumscriptions. Deflandre (1928) distinguished two subgenera, *Antarcella* and *Euarcella,* differentiated by the number of nuclei in the cell (one in the former, two or more in the latter). Recognizing 28 species in the genus, with 38 forms and varieties, Deflandre divided *Arcella* into four “sections”, according to the general morphology of their shells: Vulgares, Carinatae, Aplanatae, and Altae.

Schönborn recognized three genera in Arcellidae: *Arcella*, *Antarcella* and *Pyxidicula* (Schönborn, 1989). Since that time, the morphological delimitation of the family has been challenged by molecular data. In 2013, a multigene analysis showed that arcellinids with “secreted organic membranous…or chitinoid shells” (Arcellina *sensu* Meisterfeld, 2002) did not form a monophyletic group (Lahr et al., 2013). Subsequently, *Pyxidicula* was transferred to the Microchlamyiidae, alongside *Microchlamys, Spumochlamys* and *Microcorycia* (Porfírio-Sousa et al., 2024).

#### 3.1.2 Revision of the Arcellidae

In a major revision of the Arcellidae, González-Miguéns et al. (2022) undertook an integrative analysis of the two genera remaining in Arcellidae, *Arcella* and *Antarcella*, using the mitochondrial COI gene to test relationships within the group (González-Miguéns et al., 2022). Their molecular data supported the monophyly of Arcellidae and revealed well-resolved clades within the family. Combining these findings with morphological analysis, the authors erected the new genus *Galeripora*, which was distinguished from *Arcella* by two presumed synapomorphies: 1) the presence of pores surrounding the aperture of the shell; and 2) the presence of a “protein or organic matrix” covering at least some of the shell surface, masking its hexagonal building units. Six new species of *Galeripora* were described, and six historic species of *Arcella* were transferred to the new genus. Later work has added new species to both genera, and several more historic species have been transferred from *Arcella* to *Galeripora* on morphological grounds (Bankov, 2025; Ribeiro et al., 2023; Siemensma, 2021; Taylor et al., 2024; Useros et al., 2023).

González-Miguéns et al. (2022) examined three of Deflandre’s sections and determined that none of these groupings were monophyletic. Their molecular data indicated that the morphological features Deflandre used to define his sections were not synapomorphies but had arisen independently in different lineages by evolutionary convergence. These features of the shell were reinterpreted as environmental adaptations favoring survival in certain habitats, such as submerged vegetation, aerial moss and soil (González-Miguéns et al., 2022).

#### 3.1.3 Section Altae, Arcella leidyana and A. mitrata

Deflandre’s fourth section, Altae, was not investigated by González-Miguéns et al., 2022. In omitting the section, the authors state that Altae has not been reported from the Western Palearctic region. However, it should be noted that *Arcella mitrata* has been recorded in European samples since the turn of the last century (Stenroos, 1898; West, 1901; Cash and Hopkinson, 1905; Hoogenraad and De Groot, 1940; Siemensma, 2019). A morphotype described and illustrated by (Golemansky, 1962) from Guinee, Africa, appears to be a new species rather than *A. mitrata*. The morphotype Deflandre identified as *Arcella mitrata* var. *spectabilis* has also been found in the Netherlands (Siemensma, 2019).

Within Altae, Deflandre had placed what he called the “most aberrant forms” of *Arcella*, consisting broadly of “*Arcella mitrata* and its derivatives”: tall-shelled species whose height : diameter ratio normally exceeded 1, and could reach as much as 1.92 (Deflandre, 1928, p. 210). Deflandre included five species in the group: *Arcella jeanneli*, *A. apicata*, *A. rukiensis*, *A. leidyana* and *A. mitrata*. At that time, the first three species had been found only in Africa; since then, two of them, *A. jeanneli* and *A. apicata*, have been reported in Brazil (Gomes e Souza, 2005; Souza, 2018).

The two North American species in Altae, *A. leidyana* and *A. mitrata*, had originally been described by Leidy under the name *A. mitrata*. The latter, as Leidy conceived it, was a “very polymorphous” species (Leidy, 1879, p. 156) encompassing a wide range of morphotypes, as illustrated in Plate XXIX of his monograph. Some of the forms Leidy depicted have since been placed in separate taxa, either as varieties of *A. mitrata* or as distinct species. Deflandre, working entirely from Leidy’s text and illustrations, redescribed three of these as named varieties of *A. mitrata: v*ar. *pyriformis*, var. *gibbula* and var*. spectabilis*. Because these names were published before 1961 using the abbreviation var., they are deemed subspecific under Article 45.6.4, unless explicitly proposed as infrasubspecific. As a species-group name, it is simultaneously established at both species and subspecies rank under the Principle of Coordination (Art. 46.1), with identical authorship, date, and type. Accordingly, when treated at species rank, the valid names are: *Arcella pyriformis* Deflandre, 1928, *A. gibbula* Deflandre, 1928 and *A. spectabilis* Deflandre, 1928. Elevation to species rank requires no nomenclatural act (Siemensma, 2019).

Deflandre also established the new species *A. leidyana*, based on Figures 9, 10, and 16 from Leidy’s Plate XXIX (Deflandre, 1928). The first two of those figures represent a single individual, collected by Leidy in Absecom Pond, a “sphagnous swamp” in New Jersey; and the remaining one offers a profile view of a specimen from a bog pool in Atco, New Jersey. In both cases, the illustrations depict large, broad-based shells with nearly perpendicular sides, a peaked dorsal face and a relatively wide, shallowly invaginated aperture with a wavy margin. In his Fig. 17, Plate XXIX, Leidy presents a lateral view of a second specimen of similar size and overall shape, also collected in Absecom pond. This figure is not mentioned by Deflandre but probably depicts a member of the same taxon.^1^

Despite differences in size and morphology, the species and varieties Leidy lumped together under the name *A. mitrata* share certain distinctive features. All have crenulated apertures and relatively “tall” shells, whose ratio of height to ventral diameter exceeds 1.0. In each of the shells he illustrated, the angle of the sides, with respect to the ventral plane, is close to 90°. All specimens were found in lotic environments, often in association with *Sphagnum*.

#### 3.1.4 Arcella prismatica *and* Arcella spectabilis

In 2024, Taylor et al. described *Arcella prismatica*, a new morphospecies discovered in a bog lake in Eastern Ontario, Canada and subsequently found in the Eeyou Istchee region of western Quebec, Canada (Taylor et al., 2024).^2^ At the time of publication, molecular data were not available for this taxon, and the species was delimited by the morphology and morphometry of its shell, which is typically polyhedral, with steeply angled sides, a crenulated aperture and a nearly-flat dorsal surface. In the same investigation, the authors briefly discussed an *Arcella* from the same site which they identified as *Arcella spectabilis*. This morphotype, which often blooms alongside *Arcella mitrata* and *Arcella prismatica* in Mer Bleue bog, conforms in overall shape to the variety recognized by Deflandre as *A. mitrata* var. *spectabilis*, and to specimens of *Arcella spectabilis* collected in the Netherlands (Siemensma, 2019), although the shell is somewhat larger than previous records of the species, with a maximum diameter of 107-136 µm and a height of 109-131 µm. While several morphological characters distinguish *Arcella spectabilis* from congeners such as *A*. *prismatica*, *A. mitrata* and *A. leidyana*, the lack of molecular evidence left unanswered questions about the relatedness of these morphotypes, and their phylogenetic position within the family.

#### 3.1.5 The status of genus *Antarcella*

Deflandre, 1928, emend. Deflandre, 1953 *Antarcella*, a taxon proposed by Deflandre (1928) as a subgenus of *Arcella* and later promoted to genus level (Deflandre, 1953), was characterized by possession of a single nucleus, whereas *Arcella* had two or more. Gonzalez-Miguéns et al. (2022) invalidated *Antarcella* arguing that the uninucleate condition in some arcellid species might simply be a life stage, and that the number of nuclei in the cell has been applied inconsistently as a taxonomic character, even by Deflandre himself (González-Miguéns et al., 2022). However, it is worth mentioning that the number of nuclei is not the only distinctive character separating *Antarcella* from other Arcellidae (Meisterfeld, 2002). *Antarcella* has historically been described as possessing an ovular nucleus, in contrast to the vesicular nucleus found in *Arcella* (Collin, 1914; Penard, 1917; Meisterfeld, 2002). While vesicular nuclei typically present a single central nucleolus, sometimes accompanied by a few very small additional nucleoli, ovular nuclei contain several to many small nucleoli (Raikov, 1982). This character was not considered in the invalidation of the genus *Antarcella*; thus, further investigation incorporating nuclear morphology and nucleolar structure, potentially including nuclear staining, is needed to test the systematic hypotheses separating *Arcella* and *Antarcella* (Meisterfeld, 2002).

However, it is not only in the nucleus that *Antarcella* differs from *Arcella*. In addition to nuclear morphology, *Antarcella* differs from *Arcella* in test structure. Siemensma (2019) rediscovered *Antarcella pseudarcella* in dry mosses in Berlin (2016) and Portugal (2024). His photographs demonstrate that *A. pseudarcella* possesses pores around the aperture and, unlike *Arcella* and *Galeripora*, bears multiple xenosomes on the test—an attribute already noted by Penard (1917). Although the placement of this species within the Arcellidae cannot presently be confirmed, in our view the available morphological evidence does not call the validity of the genus *Antarcella* into question.

### 3.2 Morphological and molecular investigation of Arcellidae

To expand the morphological and COI dataset available for Arcellidae, we investigated specimens collected from Canadian peatlands (Tables 1 - 2 and Figures 2 - 4). Several study sites in Mer Bleue Bog Conservation Area and the Eeyou Istchee region provided reliable access to most of the tall-shelled forms historically recorded by Leidy (Figure 2). At two sites within the James Bay watershed (Eeyou Istchee), we found populations of *Arcella leidyana* Deflandre, 1928, a species not reliably recorded since 1879 (Table 1 and Figure 2 A-D). From these sites we also identified populations of *A. mitrata*, *A. spectabilis, and A. prismatica* (Table 1). From Mer Bleue we identified populations of *A. mitrata*, two morphotypes of *A. spectabilis* (large and small morphotypes), and *Arcella prismatica* (Table 1 and Figure 2 E-L). We also identified a population of mixotrophic specimens morphologically indistinguishable from *Galeripora artocrea* (Table 1 and Figure 3), but lacking the pores surrounding the aperture, a synapomorphic character of the genus *Galeripora*; and a morphotype characterized by a hemispherical shell and a conspicuously developed basal rim, provisionally identified as a broad-rimmed variant of *A* cf. *gibbosa*. In *Sphagnum* from a calcareous fen in Ontario, we discovered a novel terrestrial arcellid species, described here as *Galeripora purdoni* sp. nov (Table 1 and Figure 4). At another subarctic site, we collected *Galeripora dentata* Ehrenberg, 1830 (Table 1).

**Figure 2.**
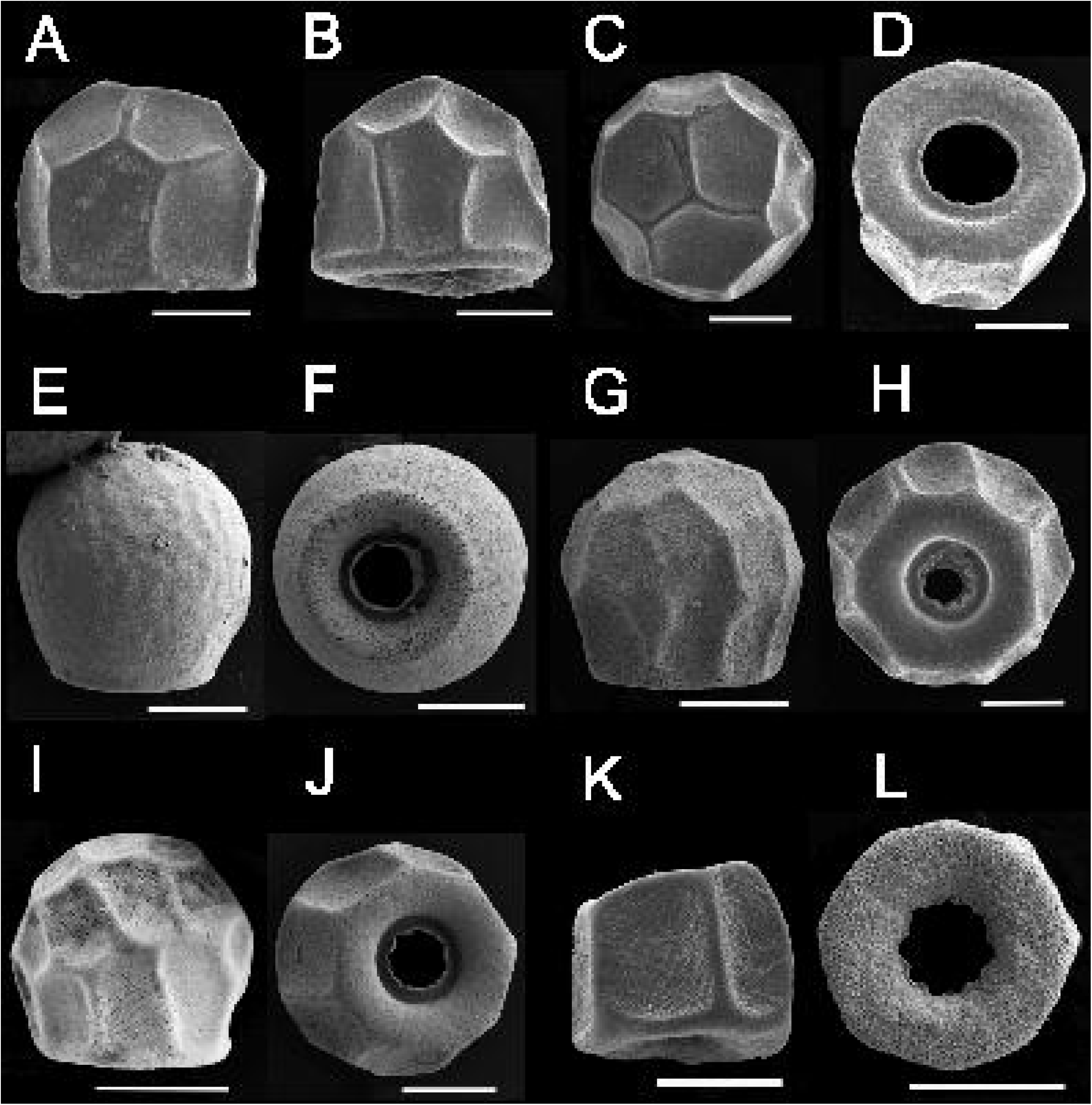
Scanning electron micrographs of *Arcella* in the North American “Altae” clade. A, B. *Arcella leidyana*, lateral views. C. *A. leidyana*, dorsal view. D. *A. leidyana*, ventral view. E,F. *Arcella mitrata*, from Mer Bleue bog. G,H. Large morphotype of *A. spectabilis*, from Mer Bleue bog. I,J. Small morphotype of *A. spectabilis*, from Mer Bleue bog. K,L. *A. prismatica*, from Mer Bleue bog. All scale bars = 50 µm.

**Figure 3.**
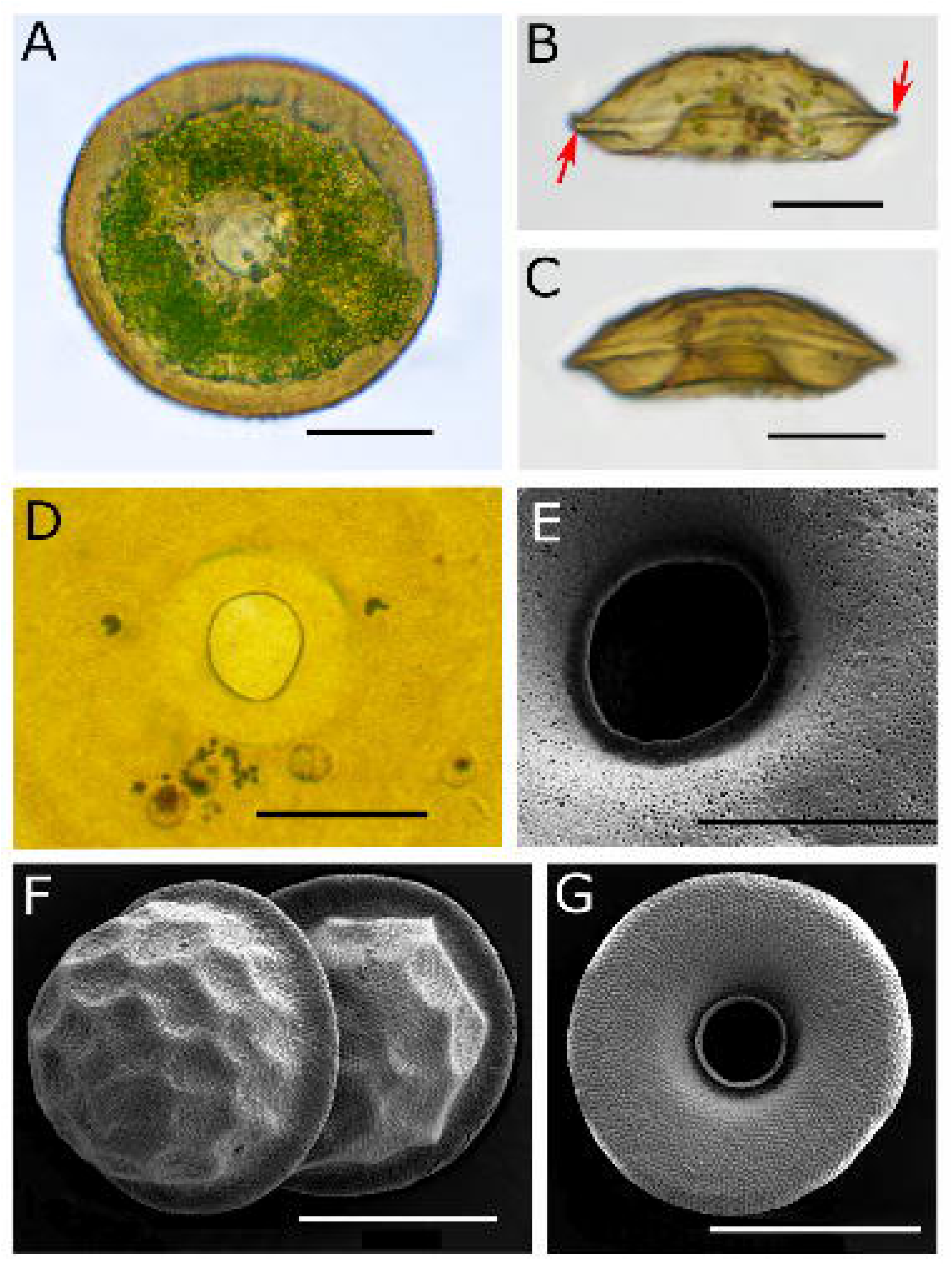
A-E. *Arcella artocrea* from Mer Bleue bog. A. Ventral view of a live specimen, showing algal endosymbionts. B,C. Lateral views of empty shells, showing the characteristic “pie shape,” deeply invaginated aperture and raised basal border (bourrelet). Arrows indicate the bourrelet. D. Ventral surface of an empty shell, showing absence of pores around the aperture. E. SEM of ventral surface, showing a field of open areoles, but no circumapertural pores. F,G. *Arcella* cf. *gibbosa* from Mer Bleue bog. F. SEM showing dorsal view of two specimens. G. SEM of ventral surface. All scale bars = 50 µm.

**Figure 4.**
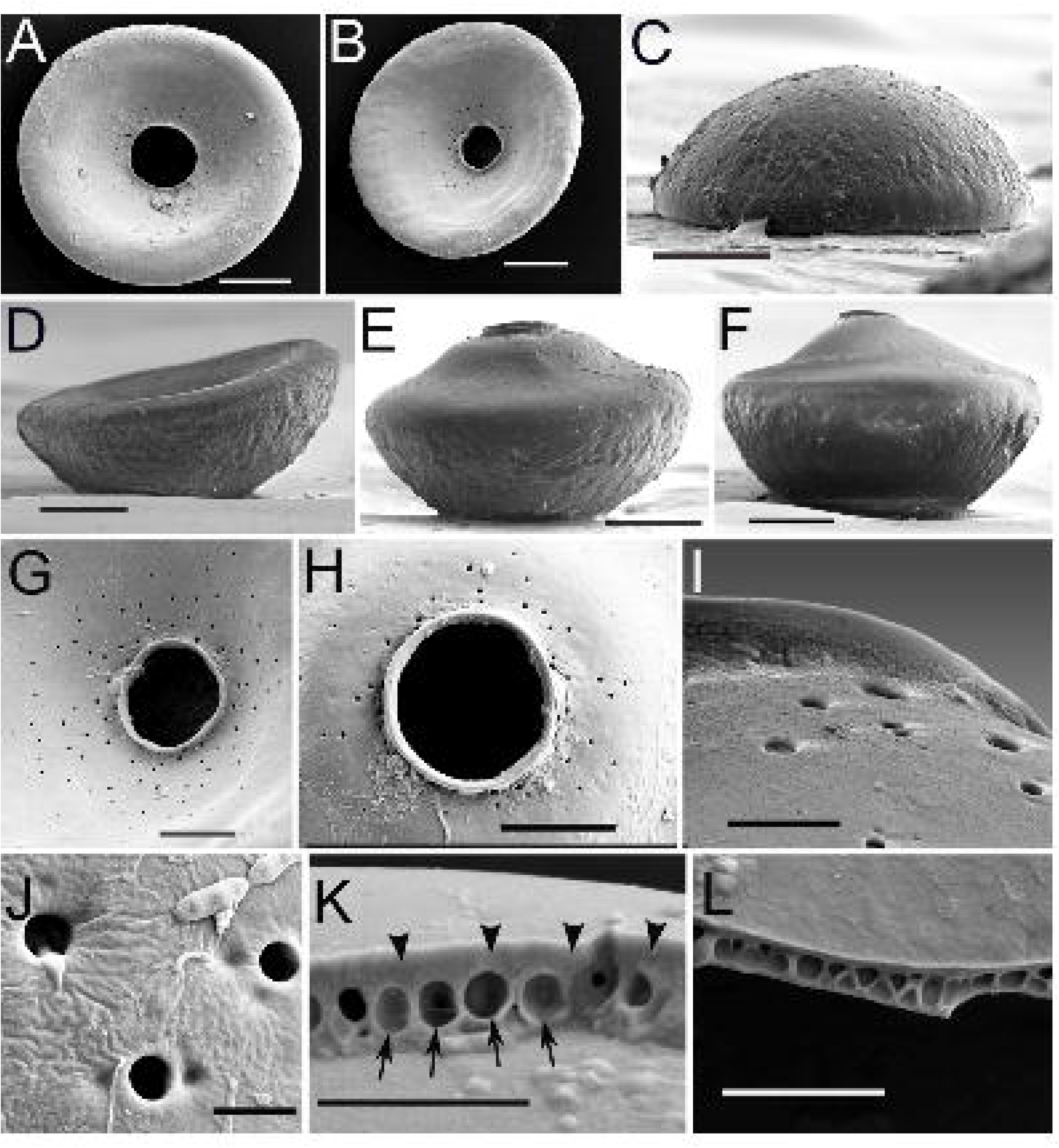
Scanning electron micrograph images of *Galeripora purdoni*. A,B. Ventral views (scale bars 50 µm). C,D. Lateral views (scale bars = 50 µm). E,F. Lateral views of encysted specimens with everted ventral surface (scale bars = 50 µm). G,H. Ventral views, showing scattered distribution of circumapertural pores (scale bars = 25 µm ). I. Lateral view of aperture, showing pores (scale bar = 5 µm). J. Close view of pores on ventral face (scale bar 2 µm). K. *Galeripora purdoni* areoles and overlying organic matrix. Black arrows indicate areoles; arrowheads indicate a layer of organic material covering the outer surface of the shell. (scale bar = 4 µm) L. A broken section of the shell, showing a loose distribution of rounded areoles.

The inferred COI maximum-likelihood (ML) phylogeny includes 64 Arcellidae taxa, comprising newly generated COI sequences, previously published data, and sequences retrieved from Arcellidae genomic-level data available through the Protist 10,000 Genomes (P10K) project (González-Miguéns et al., 2022; Ribeiro et al., 2023; Porfirio-Sousa et al., 2026). This represents a comprehensive sampling of the currently available COI data for the group. The resulting tree recovers a well-supported monophyletic clade (SH-aLRT = 98.3 / UFBoot = 100) corresponding to North American members of the *Arcella* Altae section (Figure 5). Our sampling indicates that this clade has not been previously represented in molecular datasets and suggests that its distinctive morphology may represent synapomorphies of this monophyletic clade within the genus *Arcella*. Within this clade, specimens assigned to *Arcella spectabilis* (including both large and small shell morphotypes) do not form a monophyletic group, although node support is insufficient to resolve their relationships confidently, and additional sampling is needed (Figure 5). The analysis also recovers *Arcella prismatica* as a distinct lineage within the Altae clade, corroborating its recognition as a separate species based on previous morphological evidence (Figure 5; Taylor et al., 2024). It is important to note that our dataset does not include all species assigned to the Altae section by Deflandre, and therefore it cannot be regarded as a full validation of his concept. In particular, the three tropical Altae species, *Arcella jeannelli*, *A. apicata*, and *A. rukiensis*, have not yet been sequenced, and there is no compelling reason to expect that these taxa will cluster with the North American Altae clade. Aside from differences in morphology and biogeography, the tropical species also differ in having round apertures lacking lobes or crenulations.

**Figure 5.**
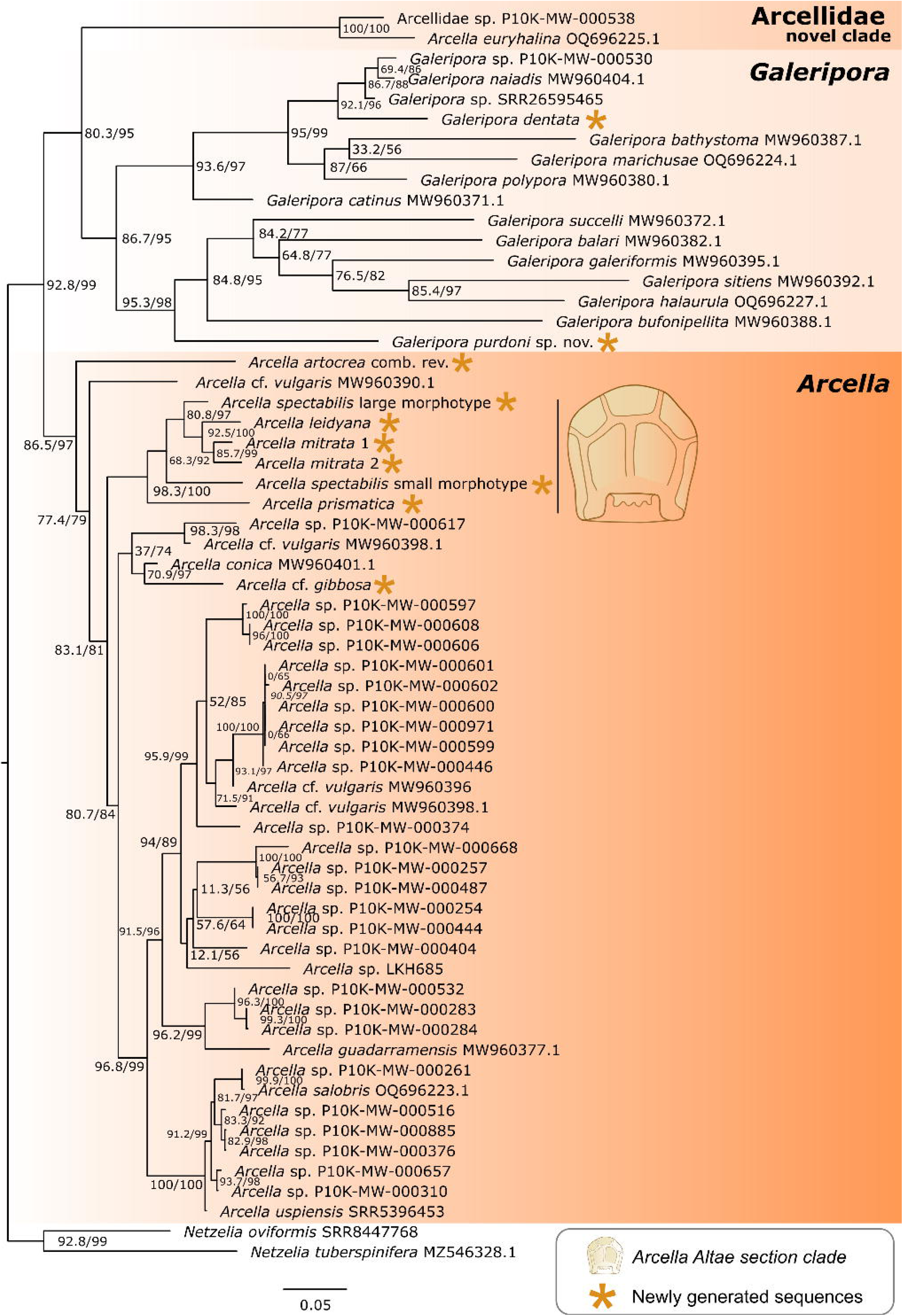
Maximum-likelihood phylogenetic tree of Arcellidae reconstructed from cytochrome c oxidase subunit I (COI) sequences. Phylogenetic reconstruction was conducted using 1,494 aligned sites in IQ-TREE v2.3.6, with ModelFinder identifying the best-fit substitution model (GTR+F+I+G4). Node support was assessed using both the Shimodaira–Hasegawa approximate likelihood ratio test (SH-aLRT) and ultrafast bootstrap (UFBoot). Support values are reported as SH-aLRT/UFBoot, with values ≥80/95 considered indicative of strong support (Guindon et al., 2010; Minh et al., 2013).

Beyond the Altae clade, the expanded sampling, including data available from the P10K project, reveals several additional well-supported clades within *Arcella*. These lineages likely differ in genomic, genotypic, and ecological traits, as suggested in previous studies. Some are placed in lower-supported clades, including the newly sampled *A*. cf. *gibbosa*, suggesting potentially undersampled regions of the *Arcella* phylogeny. A comprehensive integrative framework combining genomic, morphological, and phylogenetic data will be necessary to further investigate their diversity and evolutionary relationships.

The mixotrophic population identified as *Galeripora artocrea* corresponds closely to the morphotype described by Leidy in 1876 and later illustrated in his 1879 monograph (see Leidy’s figs. 1, 2, 9; Pl. XXX). In our phylogenetic analyses, this population does not branch among *Galeripora* species but instead clusters with *Arcella* in a basal position (Figure 5). Light and scanning electron microscopy corroborate this placement: the morphotype lacks pores around the shell aperture, a key diagnostic feature of *Galeripora* (Figure 3). These results indicate that *Galeripora artocrea*, as currently circumscribed, is polyphyletic. Because our specimens conform to the typical form described by Leidy and recognized by Deflandre as the nominotypical expression of *Arcella artocrea*, we return the species to its original genus as *Arcella artocrea* Leidy, 1876 comb. rev. According to Art. 45.6.4 and Art. 46.1 of ICZN, the correct name for *Galeripora artocrea* var. *pseudocatinus* (Deflandre, 1928) is *Galeripora pseudocatinus* (Deflandre, 1928).

Within *Galeripora*, the newly generated *G. dentata* sequence branches close to previously sampled *Galeripora* from freshwater habitats. In addition, the newly sampled *Galeripora purdoni* sp. nov. from soil, exhibiting distinctive morphological characteristics compared to previously described species, branches with the clade that includes *G. sitiens*, *G. halaurula*, *G. bufonipellita*, *G. balari* and *G. galeriformis*, all of which have been identified as primarily terrestrial species (Useros et al., 2023).

Our expanded COI sampling, including data from the P10K project, identified the Arcellidae isolate P10K-MW-000538, which branches with full support (SH-aLRT = 100 / UFBoot = 100) as sister to the recently described *Arcella euryhalina* (Figure 5). In the original description, although COI data did not corroborate this placement, *A. euryhalina* was assigned to *Arcella* due to the absence of pores surrounding the shell aperture, as the presence of pores constitutes the key diagnostic character of *Galeripora*. In our COI phylogeny, *A. euryhalina* and the P10K-MW-000538 isolate are recovered outside the *Arcella* clade and closer to *Galeripora*, although this relationship is not fully supported (SH-aLRT = 80.3 / UFBoot = 93; Figure 5). Since P10K isolates are represented by genome-scale data, we further investigated P10K-MW-000538 using phylogenomic analyses. The phylogenomic reconstruction robustly places P10K-MW-000538 as a basal lineage within Arcellidae, distinct from both the *Arcella* and *Galeripora* clades (Figure 6). Under the current taxonomy of the group, these results suggest that *Arcella euryhalina* and P10K-MW-000538 may represent a third genus within Arcellidae. However, due to the lack of comprehensive morphological data for these isolates, we refrain from formal taxonomic action. Both originate from high-salinity environments, and together they provide a reference for future sampling efforts targeting this lineage, which expands the known diversity of Arcellidae and is supported by both COI and multi-gene phylogenetic reconstructions. The concatenated analysis of the COI and phylogenomic datasets corroborated the topologies recovered from the individual COI and phylogenomic datasets (Supplementary Fig. 2).

**Figure 6.**
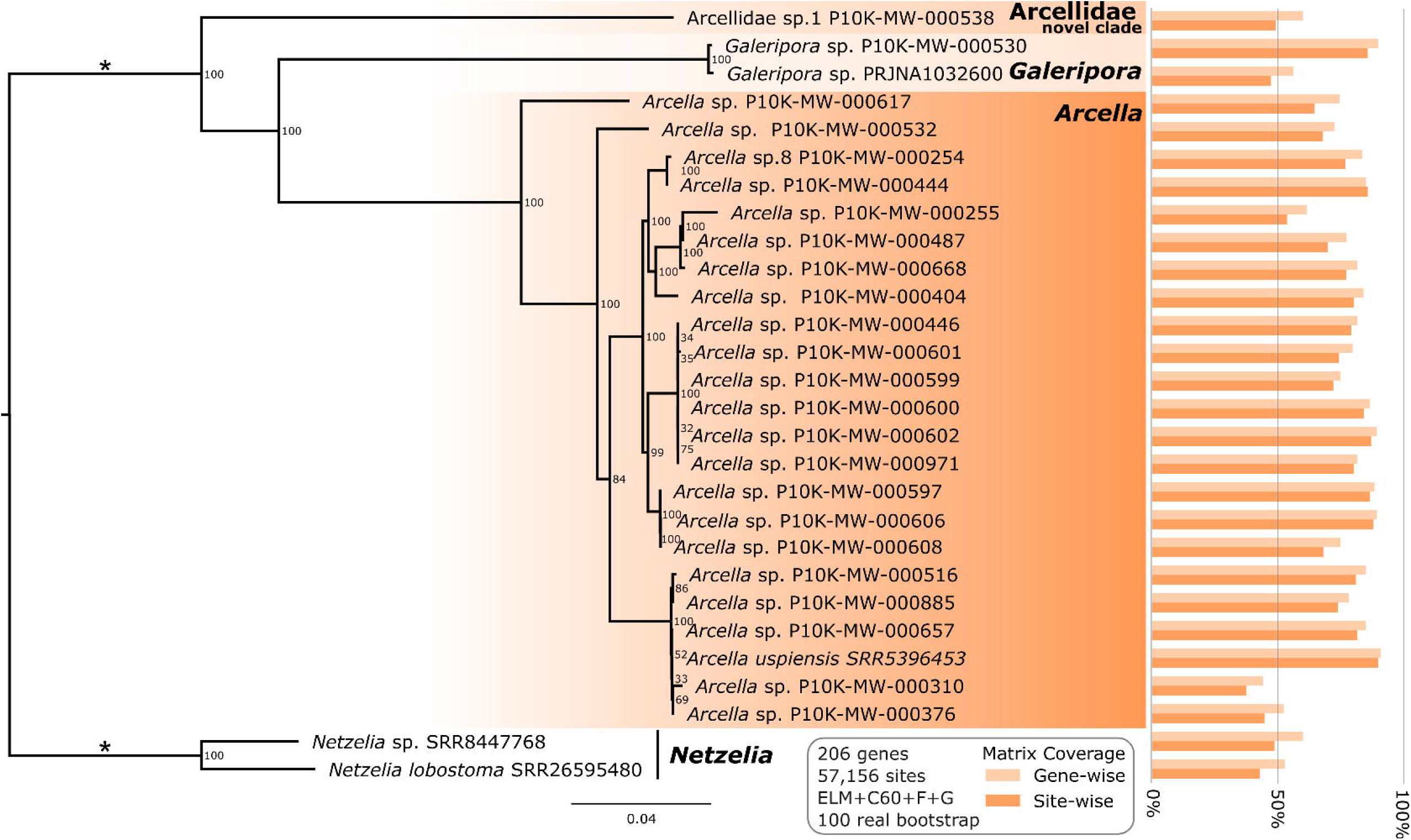
Phylogenomic tree of Arcellidae. Phylogeny of Arcellidae and *Netzelia* based on 206 genes (57,156 amino acid sites). The tree was initially built using IQ-TREE v. 3.0.1 under the ELM+C20+F+G site heterogeneous model of evolution and further used to infer a posterior means site frequency model using the ML model ELM+C60+F+G model of evolution in IQ-TREE v. 3.0.1, collecting posterior mean site frequencies (PMSF) inferred from the dataset. Topological support was assessed by 100 non-parametric real bootstrap replicates under the ELM+C60+F+G+PMSF model of evolution in IQ-TREE3. The length of the branches indicated by a * has been reduced by 50%. The bar plots on the right side of the figure show the percentage completeness of the dataset available for each taxon, based on the total number of genes and sites in our phylogenomic tree matrix.

The results presented here highlight both the strengths and current limitations of integrative approaches to Arcellidae systematics. The monophyly of North American Altae illuminates a hypothesis that had remained untested since Deflandre’s original proposal, and the concordance between molecular and morphological evidence in *A. prismatica*, *A. leidyana*, *A. artocrea* and *G. purdoni* sp. nov. corroborates that COI barcoding, when combined with careful morphological characterization, provides a robust framework for species delimitation. The identification of a newly sampled lineage within Arcellidae represented by *A. euryhalina* and corroborated by the phylogenomic analysis of the P10K-MW-000538 isolate further underscores the power of genome-scale data to reveal deep evolutionary structure that single-marker approaches may fail to resolve. Nevertheless, important uncertainties remain, including the poorly resolved internal relationships within Altae, the apparent polyphyly of *A. spectabilis*, and the ambiguous placement of the *A. euryhalina* lineage under COI data, all of which reflect both marker limitations and incomplete taxon sampling, particularly for tropical, cryptic, and high-salinity lineages. The occurrence of *G. purdoni* sp. nov. in a calcareous fen also suggests that ecologically marginal habitats may harbor unrecognized diversity shaped by substrate chemistry and local environmental filtering. Consequently, a comprehensive exploration of Arcellidae diversity will require expanded geographic and ecological sampling, multi-locus and phylogenomic datasets beyond current coverage, and renewed examination of historically described taxa, with Canadian boreal and subarctic peatlands continuing to provide particularly valuable material for this integrative framework. Importantly, the combined morphological and molecular evidence presented here supports three taxonomic acts, namely the description of *Galeripora purdoni* sp. nov., and the redescription of *A. leidyana* and *A. artocrea* comb. rev.

### 3.3 Taxonomic actions

#### 3.3.1 Description of *Galeripora purdoni* sp. nov

Order Arcellinida Kent, 1880

Infraorder Sphaerothecina Kosakyan et al., 2016

Family Arcellidae Ehrenberg, 1830

Genus *Galeripora* González-Miguéns et al., 2022

Galeripora purdoni sp. nov.

##### ZooBank registration number of the present work

(LSID to follow)

##### Zoobank registration of the new species

(LSID to follow)

##### Examined material

56 specimens with LM, 23 specimens with SEM, (Purdon Conservation Area, Lanark, Ontario, Canada)

##### Diagnosis

Shell irregularly circular in apical view. Dorsal surface domed and densely pitted with small depressions, becoming smooth toward the margins. Ventral surface smooth and concave in active specimens, bearing numerous pores (>30, sometimes >100) that are loosely scattered across the surface rather than confined to the circumapertural region. Pores are larger and more abundant near the aperture, becoming smaller and sparser toward the shell margin. Aperture central, circular to ovoid, deeply invaginated, with a slightly everted collar. All exposed surfaces thinly coated with organic material that conceals the areolar building units. During encystment, the ventral surface everts and forms a domed or conical protuberance over the spherical resting cyst.

##### Etymology

The species epithet honors Joseph “Joe” Purdon (1914-1982), a boatbuilder and self-taught conservationist who spent nearly fifty years nurturing a rare population of Showy Lady’s Slipper orchids on his property in eastern Ontario. After his death in 1982, stewardship of the site passed to the Mississippi Valley Conservation Authority, which continues to maintain this sensitive wetland as the Purdon Conservation Area (White, 1988). Purdon’s devotion to one vulnerable species enabled the permanent protection of all the organisms that share this unique habitat.

##### Type specimens (Holotype and paratypes)

Permanent LM slides and SEM stubs deposited at the Canadian Museum of Nature (accession numbers to follow)

##### Type locality

Purdon Conservation Area, Lanark, Ontario (44°59’32.5“N 76°32’44.6”W), a calcareous woodland fen, among hummocks of *Sphagnum* and embedded forest litter.

##### Description of the type material

All observed specimens had conspicuous discoid shells, 179–254 μm in diameter (mean 212 μm, n=56), varying in color from pale yellow to reddish brown. Shells of active specimens were subcircular and often slightly asymmetrical. Vacant shells were easily crumpled and could appear ellipsoid, obovate, or even convexly triangular. The aperture was small, 77–102 μm in diameter (mean 83 μm, n=56), and imperfectly circular, bordered by a narrow collar of organic cement. The apertural invagination was 34–43 μm high (mean 39 μm, n=56) with a ratio of aperture to shell diameter of 0.18–0.24 (mean 0.21).

The aboral (dorsal) surface was domed, 77–102 μm high (mean 83 μm), and marked by shallow indentations, becoming smooth at the basal margin. The ventral surface was smooth and concave in active specimens, giving the shell a bowl like appearance. In encysted specimens, the ventral surface was displaced outwards, becoming more or less mammiform.

Pores on the ventral surface varied in size, generally being larger around the aperture and very small at the shell margins. All exposed surfaces were coated with secreted material that conceals the areoles, except where breakage revealed them in cross section. Areoles measured approximately 750–950 nm in diameter. See Supplementary Table S5 for basic morphometrics of the population from Purdon Fen.

##### Differential diagnosis

*Galeripora purdoni* is distinguished from congeners by molecular markers in the mitochondrial COI gene and by the exceptionally large number of pores scattered across its ventral surface. In ventral view, it can be differentiated from *G. discoides*, *G. marichusae*, *G. megastoma*, *G. polypora*, and *G. scutelliformis* by the small size of its aperture relative to the overall shell diameter. In lateral view, it differs from all varieties of *A. artocrea* and *G. arenaria* by the absence of a distinct border or basal rim on the shell margin (and from the latter also by its much larger size). The dorsal (apical) surface of the shell is distinguished from that of *G. catinus* and *G. balari* by the lack of ridges, folds, or facets. *Galeripora succelli* has a similar size and profile, but its test is described as flattened at the margins (i.e. with a “keel”), and its pores are restricted to a ring around the aperture. Other species in the genus—*G. bathystoma*, *G. bufonipellita*, *G. galeriformis*, *G. halaurula*, *G. naiadis*, and *G. sitiens*—may share individual characters with *G. purdoni*, but all are much smaller in size.

One taxon, described by Decloitre (1972) as *Arcella arenaria* var. *irregularis*, has a loose scattering of variably sized pores around the aperture, somewhat like that of *G. purdoni*. However, Decloitre’s variety is much smaller (70-80 μm, less than half the size of *G. purdoni*), and was reported from lichen on olive trees.

See Supplementary Table S6 for a comparison of basic morphometrics and diagnostic characters of all described species and varieties in the genus.

#### 3.3.2 Emended diagnosis of *Arcella leidyana*

Order Arcellinida Kent, 1880

Infraorder Sphaerothecina Kosakyan et al., 2016

Family Arcellidae Ehrenberg, 1830

*Arcella* Ehrenberg, 1830

*Arcella leidyana* Deflandre, 1928 Zoobank registration

(LSID to follow)

##### Examined material

47specimens with LM and 34 specimens with SEM (Chisabi, Eeyou Istchee, Canada)

##### Emended diagnosis

Shell polyhedral, with 6–10 ribs (usually 8); dorsal surface faceted, forming a peak, gable or angular dome. In apical view, shell is polygonal or nearly round. In lateral view, approximately house-shaped, aborally peaked with nearly parallel sides almost perpendicular to the ventral plane (mean angle 89°). Aperture notably wide and shallowly invaginated; central, approximately circular, with subtle crenulations forming a wavy margin around the raised lip. The ratio of aperture to ventral diameter ranges from .38 to .47 (mean .42).

##### Voucher specimens (topotypes)

Permanent LM slides and SEM stubs deposited at the Canadian Museum of Nature (Accession number to follow)

##### Description of the observed specimens

The shells, composed of hexagonal areoles, were nearly colorless when newly formed, gradually shifting to yellow, amber, and eventually brown. Older specimens frequently darkened to a greyish indigo color, presumably through complexation of iron within the organic matrix of the test (cf. Deflandre, 1928, p. 173). In old, vacant shells—particularly those that had dried—the outer areolar faces often collapsed, leaving a field of hollow depressions across the test surface. The maximum diameter of all 47 observed shells was 116–153 μm, mean 134 μm. The shell height was 103–134 μm, mean 116 μm. The aperture diameter was 41–68 μm, mean 58 μm, and the aperture height 10–20 μm, mean 14 μm.

In ventral view, polygonal shells lacked a “concentric circle” around the aperture, a feature that marks the presence of a deep buccal tube in morphotypes assigned by Deflandre to *Arcella mitrata* (Deflandre, 1928, pp. 270–71). The aperture itself was conspicuously wide, spanning 42–47% of the ventral diameter (a ratio nearly twice as large as that in *Arcella spectabilis*). In profile, the apex of the shell was always raised above the perpendicular side-walls, forming a pyramidal peak, faceted dome or gabled ridge. Pseudopods were lobose and often numerous, but typically quite short, rarely extending beyond the margins of the shell. See Supplementary Table S7 for basic morphometrics of the population from Chisasibi.

##### Differential diagnosis

*Arcella leidyana* is set apart from congeners by molecular markers in the mitochondrial COI gene. Morphologically, it differs from *A. spectabilis* in its profile, with nearly parallel lateral walls rising to a peaked apex, and in the shallow invagination of its very broad aperture. It is readily distinguished from *A. prismatica* by the dorsal (apical) surface of the test, which is never flat, and by the aperture, which lacks strongly defined lobes. Its overall form resembles that of *A. conica* Playfair, 1918, described by its author as having “the shape of a marquee tent”; however, *A. conica* possesses a compact, smoothly rounded aperture without crenulations, and is a much smaller species (diameter 68–100 µm).

#### 3.3.3 Emended diagnosis of *Arcella artocrea*

Order Arcellinida Kent, 1880

Infraorder Sphaerothecina Kosakyan et al., 2016 Family Arcellidae Ehrenberg, 1830

*Arcella* Ehrenberg, 1830

Arcella artocrea Leidy, 1876

##### Zoobank registration

(LSID to follow)

##### Examined material

84 specimens (Mer Bleue, Ottawa); 48 specimens (Chisasibi, Quebec)

##### Emended diagnosis

Shell approximately circular in ventral view, sometimes with local indentations and irregularities. Ventral surface smoothly convex, forming a raised border one-third to one-half of total shell height. Test border well defined, often with a thickened projection on the rim (bourrelet). Dorsal (aboral) surface convexly domed, bearing pits or mammilations. A circular depression is sometimes present around the lower perimeter of the dorsal dome, internal to the bourrelet. In lateral view, the raised basal border forms a horizontal ledge projecting on each side of the dorsal dome. Aperture invaginated to approximately one half of shell height, with an everted lip. Cytoplasm with green algal endosymbionts.

##### Voucher specimens (topotypes)

Permanent LM slides deposited at the Canadian Museum of Nature (Accession number to follow)

##### Description of the observed specimens

Healthy, living specimens from locations in Ottawa and Chisasibi were abundantly filled with green algae, conspicuous at low magnification as an irregular dark mass within the test. Shells were light amber to golden, with minimal darkening in older individuals. Pseudopods were narrow and tubular, often long, not numerous. The shell diameter of specimens from Mer Bleue was 130–159 μm (n=84); the height was 41–56 μm (n=15) and the diameter of the aperture 19–33 μm, with ratio of aperture to diameter 0.13–0.23 (n=66). The shell diameter of specimens from Chisasibi was 141-178 μm (n=48); the diameter of the aperture was 21-31 μm, with ratio of aperture to diameter 0.12-0.2 (n=28).

##### Differential diagnosis

Distinguished from *Galeripora arenaria* and *G. pseudocatinus* comb. nov., by its persistent green algal endosymbionts, the absence of circumapertural pores, and the presence, in most specimens, of a tubular ridge (bourrelet) around the basal rim or a circular depression between the raised border of the shell and the dorsal dome.

##### Remarks

Leidy described *A. artocrea* as a “singular pie-shaped *Arcella* with a bright green sarcode,” emphasizing that algae are a stable component of the cytoplasm: “Entosarc loaded with chlorophyll balls which appear to be an element of structure” (Leidy, 1876). In his original account he did not mention circumapertural pores, and none are shown in his first illustrations of the species (Leidy, 1879: Pl. XXX, figs. 1, 2, 9). In contrast, figs. 3–8 of the same plate depict a second morphotype lacking a thickened basal rim and bearing approximately 20 pores around the aperture, which Leidy interpreted as “minute tubercles.” Noting these differences, Deflandre (1928) separated the pore-bearing form as *Arcella artocrea* var. *pseudocatinus*. In Mer Bleue, the typical form of *Arcella artocrea*—pie-shaped, with a bourrelet—consistently contains abundant algal endosymbionts, a feature not known from any other known species of Arcellidae.

#### 3.3.4 *Galeripora pseudocatinus* (Deflandre, 1928) comb. nov

##### Zoobank registration

(LSID to follow)

The name *Arcella artocrea* var. *pseudocatinus* Deflandre, 1928 is an available infrasubspecific name published before 1961 and therefore already constitutes a species-group name under the ICZN. Elevation to species rank requires no nomenclatural act. Only the transfer to *Galeripora* constitutes a nomenclatural act, expressed as: *Galeripora pseudocatinus* (Deflandre, 1928), comb. nov.

The results reported here advance the integrative taxonomy of Arcellidae on several fronts but also make clear how much remains to be done. The monophyly of the North American Altae clade, well-supported in both our COI and phylogenomic reconstructions, corroborates a hypothesis that had stood untested for nearly a century, yet the absence of sequence data for the three tropical Altae species means that the section as Deflandre conceived it cannot yet be evaluated as a whole. Similarly, molecular data are needed for tall-shelled species described since Deflandre—in particular, *Arcella peruviana* Reczuga et al., 2013 and *A. nordestina* Vucetich, 1973 (Reczuga et al., 2015; Vucetich, 1973).

The apparent polyphyly of *A. spectabilis* is similarly unresolved and reflects a recurring difficulty in Arcellidae systematics: morphologically coherent and visually distinctive forms that do not map cleanly onto molecular lineages, leaving open the possibility of either overlooked species boundaries or genuine intraspecific variation with a genetic component. Biogeographic patterns within the family also warrant closer scrutiny. Several of the morphotypes documented here, including *A. mitrata* and *A. spectabilis* are abundant and ecologically conspicuous in eastern North American peatlands yet appear to be scarce or unreported across much of Europe, a pattern for which neither dispersal limitation nor environmental filtering offers an obvious explanation at present. *Arcella prismatica*, despite now being recorded from multiple sites across eastern North America, has not been found outside the continent. Whether this reflects genuine endemism, insufficient sampling elsewhere, or morphological crypsis under a different name in other floras, is a question that an expanded barcoding effort in European and Asian peatlands could begin to answer.

The rediscovery of the mixotrophic form of *Arcella artocrea* raises similar issues. This morphotype has apparently not been recorded since Leidy’s description of 1879, in which it is combined with *Galeripora pseudocatinus*. Although “*Arcella artocrea*” is frequently reported in ecological and paleontological studies from sites in Europe, Asia and the Americas, we have found no specific mention of other populations containing algal endosymbionts. After Leidy, new published records of “*Arcella artocrea*” and “*Galeripora artocrea*” that include morphological data have described or depicted heterotrophic specimens bearing a ring of pores around the aperture and must therefore be referred to *Galeripora*. The polyphyly of *Galeripora artocrea*, as previously circumscribed, presents an interesting case of evolutionary convergence. Both morphotypes are associated with wet or aquatic conditions, so it is possible that their shared morphology reflects common environmental pressures.

The identification of a third lineage of Arcellidae through phylogenomic analysis of P10K isolates, associated with high-salinity environments and morphologically uncharacterized, points to a different kind of gap: organisms that fall outside the ecological contexts in which protistologists have traditionally searched for Arcellidae, and that are consequently invisible to conventional sampling strategies. Together, these findings reinforce the view that the known diversity of Arcellidae represents a fraction of what exists, and that progress toward a comprehensive classification of the family will require not only more sequences, but a broader conception of where its members are likely to be found.

## Acknowledgements

We thank Enrique Lara, Científico Titular, Real Jardín Botánico de Madrid, for advice on DNA extraction and amplification, and Glenn Poirier, Senior Research Assistant, Mineralogy, and Electron Microprobe Laboratory Manager, for assistance with electron microscopy. We are grateful to the National Capital Commission for sampling permits at Mer Bleue Bog, and to the Mississippi Valley Conservation Authority for permission to use data and materials from Purdon Conservation Area. Special thanks to Ernest and Gary Webb of Chisasibi for their help with field sampling on Fort George Island. The authors acknowledge the High Performance Computing Center (HPCC) at Texas Tech University for providing computational resources that have contributed to the research results reported within this paper. URL: http://www.hpcc.ttu.edu. This work was supported by the Gordon and Betty Moore Foundation, GBMF13832 and Grant DOI: 10.37807/GBMF13832 provided to A.K.T.

## Footnotes

1 Another large, faceted morphotype is shown in figure 11 from the same plate. However, this shell has a deeply invaginated aperture, as well as a distinct constriction above the base. Because of these features it is unlikely to be *A. leidyana*. It resembles the large morphotype of *Arcella spectabilis* that is common in samples from Mer Bleue bog and sites in Eeyou Istchee.

2 The species has since turned up at three sites in the eastern United States, in Massachusetts, New Jersey and South Carolina (iNaturalist community. 2026. Observations of *Arcella prismatica* from the United States, observed between May 6, 2025 and Nov. 8, 2025. Exported from https://www.inaturalist.org <u>on 14 May 2026.)</u>

