## Supplementary Material for "Integrative morphology and phylogenetics of Arcellidae (Amoebozoa:Arcellinida), with redescription of *Arcella leidyana* and *Arcella artocrea* and description of *Galeripora purdoni* sp. nov"

Authors: Bruce D. S. Taylora,1,*, Alfredo L. Porfirio-Sousab,*, Robert E. Jonesb, Carl Seaquistb, Ferry J. Siemensma c, Elias Taylor d, Alexander K. Ticeb,1

[a] Canadian Museum of Nature, PO Box 3443, Station D, Ottawa, ON K1P 6P4

b Department of Biological Sciences, Texas Tech University, Lubbock, TX, USA

c Julianaweg 10, Kortenhoef, 1241VW, Netherlands

d University of Guelph, Advanced Analysis Centre: Genomics Facility, Guelph, ON, N1G-2W1, Canada

* Contributed equally

**This file includes:**

Supplementary Tables S1 – S7 legends

Supplementary Figs. 1 - 7

**Supplementary table legends**

**Supplementary Table S1.** Cytochrome C oxidase subunit I (COI) dataset (Figure 5), sequences ID, P10K and NCBI accession number.

**Supplementary Table S2.** Phylogenomic matrix data information. a. Phylogenomic Matrix (Figure 6), Gene Compostition and Indices for Concatenated Matrix. b. Phylogenomic Matrix (Figure 6), Gene Occupancy Data for Each Taxon.

**Supplementary Table S3.** Arcellidae phylogenomic supermatrix (Figure 6).

**Supplementary Table S4.** Cytochrome C oxidase subunit I (COI) dataset (Supplementary Fig. 1.), sequences ID, P10K and NCBI accession number, including all newly generated COI sequences.

**Supplementary Table S5.** Basic morphometrics of *Galeripora purdoni* from Purdon Fen.

**Supplementary Table S6.** Comparison of basic morphometrics and diagnostic characters of described species and varieties in the genus *Galeripora*.

**Supplementary Table S7.** Basic morphometrics of *Arcella leidyana* from Chisasibi.

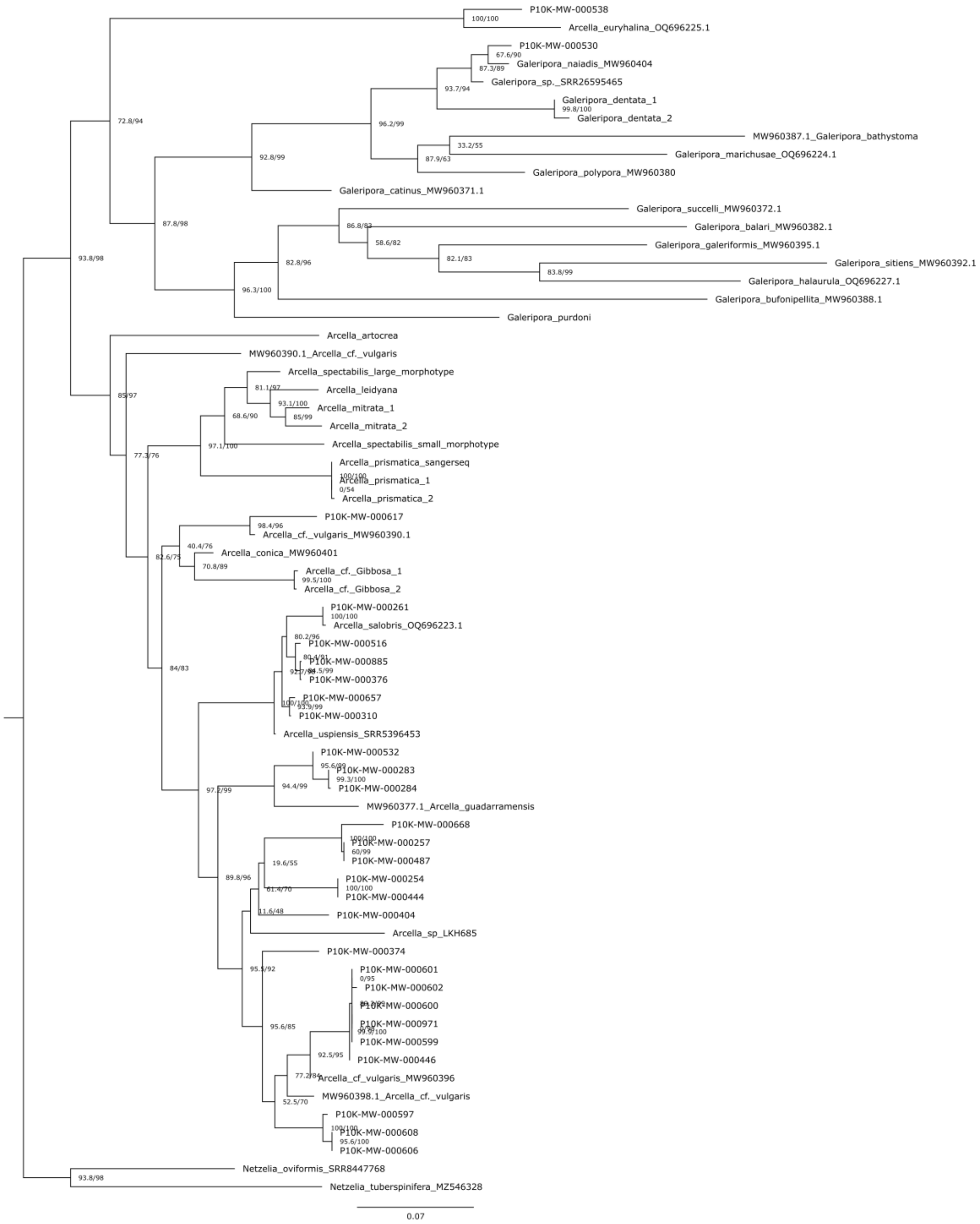


**Supplementary Fig. 1**. Maximum-likelihood phylogenetic tree of Arcellidae reconstructed from cytochrome c oxidase subunit I (COI) sequences, including all newly generated COI sequences (Table S4). The phylogenetic reconstruction was conducted using 1,485 aligned sites in IQ-TREE v2.3.6, with ModelFinder identifying the best-fit substitution model (GTR+F+I+G4). Node support was assessed using the Shimodaira–Hasegawa approximate likelihood ratio test (SH-aLRT) and ultrafast bootstrap (UFBoot). Support values are reported as SH-aLRT/UFBoot, with values ≥80%/95%, respectively, considered indicative of strong support (Guindon et al., 2010; Minh et al., 2013).

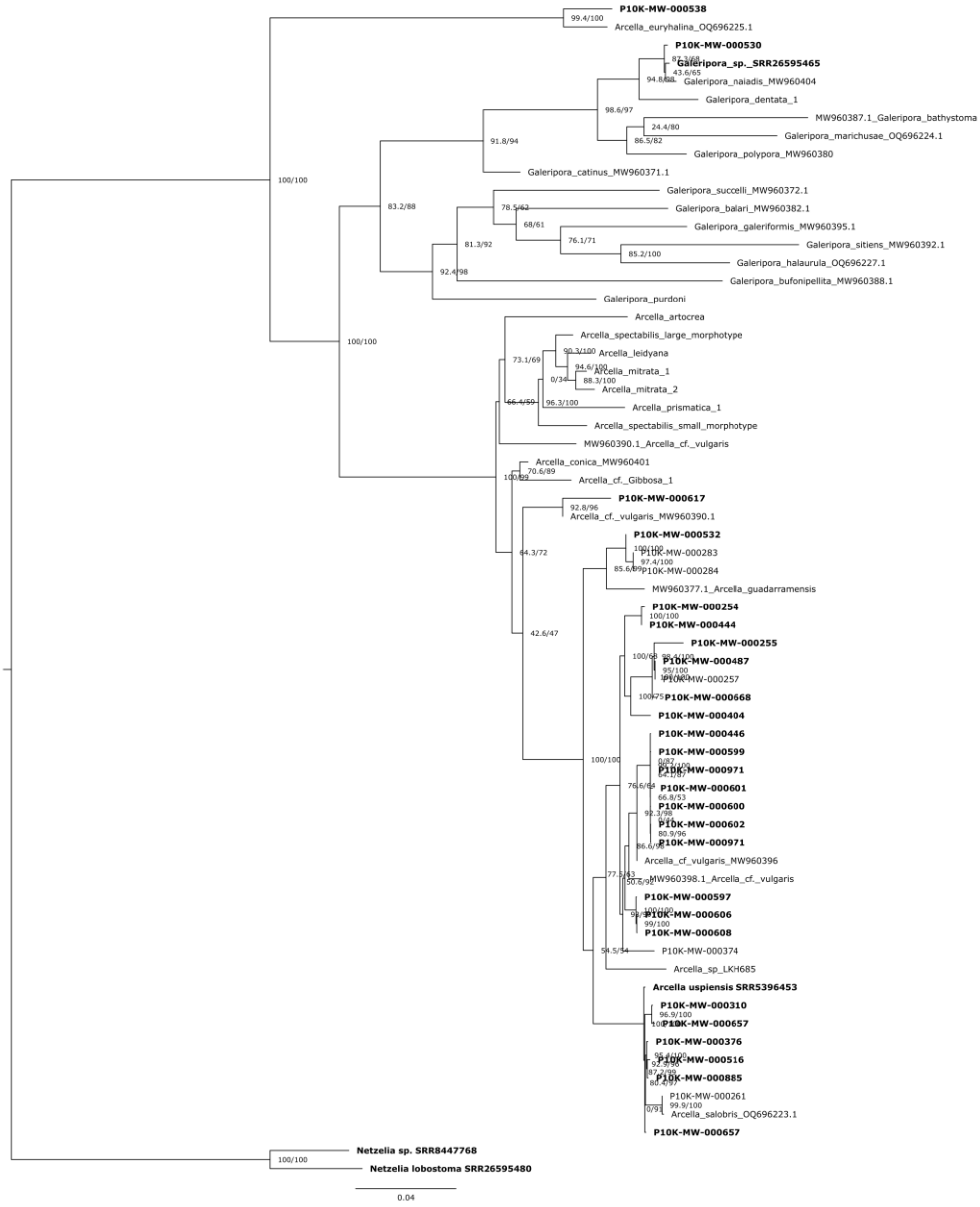


**Supplementary Fig. 2.** Maximum-likelihood phylogenetic tree of Arcellidae reconstructed from a concatenated dataset combining the cytochrome c oxidase subunit I (COI) dataset (Table S1) and the PhyloFisher phylogenomic dataset (Table S3). The phylogenetic reconstruction was conducted using 58,644 aligned sites in IQ-TREE v3.1.1, with separate substitution models applied to each partition: ELM+C20+F+G for the PhyloFisher phylogenomic dataset (57,156 sites) and GTR+F+I+G4 for the COI dataset (1,488 sites). Node support was assessed using the Shimodaira–Hasegawa approximate likelihood ratio test (SH-aLRT) and ultrafast bootstrap (UFBoot), each with 1,000 replicates. Support values are reported as SH-aLRT/UFBoot, with values ≥80%/95%, respectively, considered indicative of strong support (Guindon et al., 2010;

Minh et al., 2013). Bold terminal labels indicate taxa for which PhyloFisher phylogenomic data were available and included in the concatenated dataset.

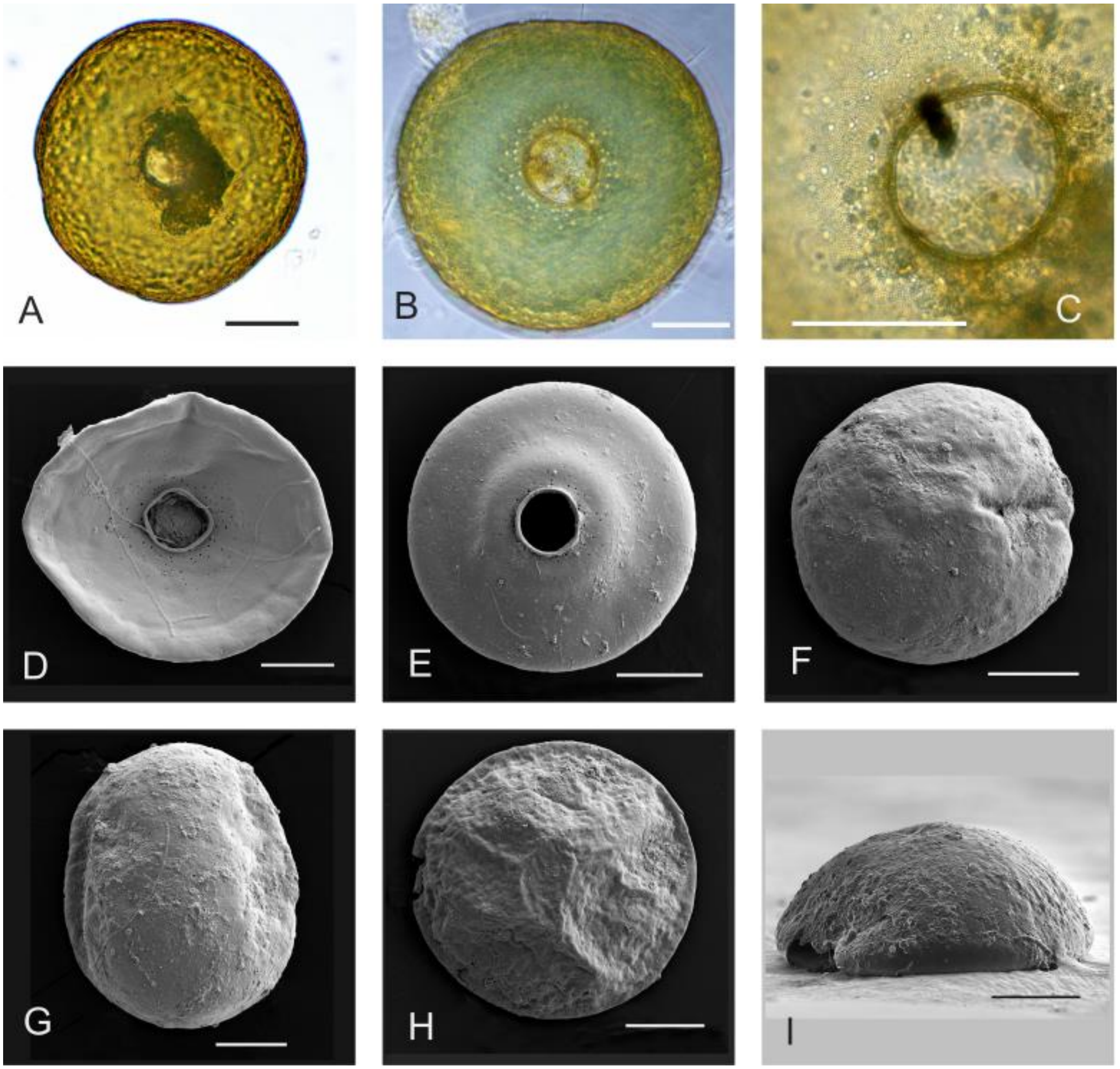


**Supplementary Fig. 3.** A-I. *Galeripora Purdoni*, from Purdon Fen, Lanark, Ontario, Canada. A,B. Ventral views; compound light microscope, 400×. C. Ventral; LM, 1,000×. D. Ventral; scanning electron microscope, 694×. E. Ventral, encysted specimen; SEM, 758×. F. Dorsal; SEM, 824×. G. Dorsal view of an oval individual; SEM, 824×. H. Dorsal; SEM, 733×. I. Profile; SEM, 1,287×.  All scale bars = 50 μm.

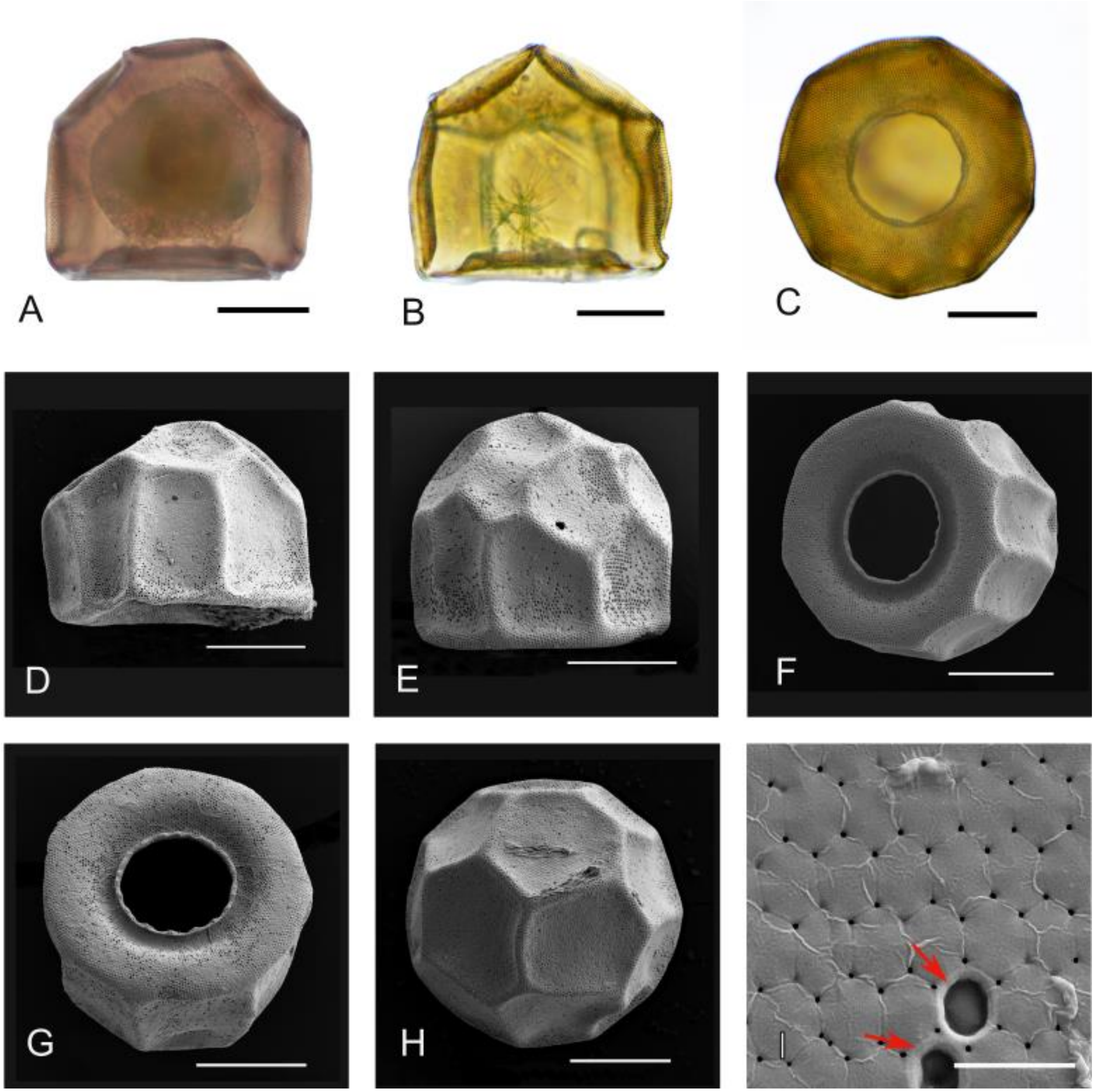


**Supplementary Fig. 4.** A-I. *Arcella Leidyana*, from Chisasibi, Quebec, Canada. A,B. Profile views; LM, 400×. C. Ventral; LM, 400×. D,E. Profiles; SEM, 1,128× and 1,200×. F,G. Ventral; SEM, 1,045× and 1,000×. H. Dorsal; SEM, 908×. I. Hexagonal building units of the shell; red arrows show two collapsed areoles; SEM, 15,000×. All scale bars = 50 µm, except Fig. 4.I., where the scale bar = 5 µm.

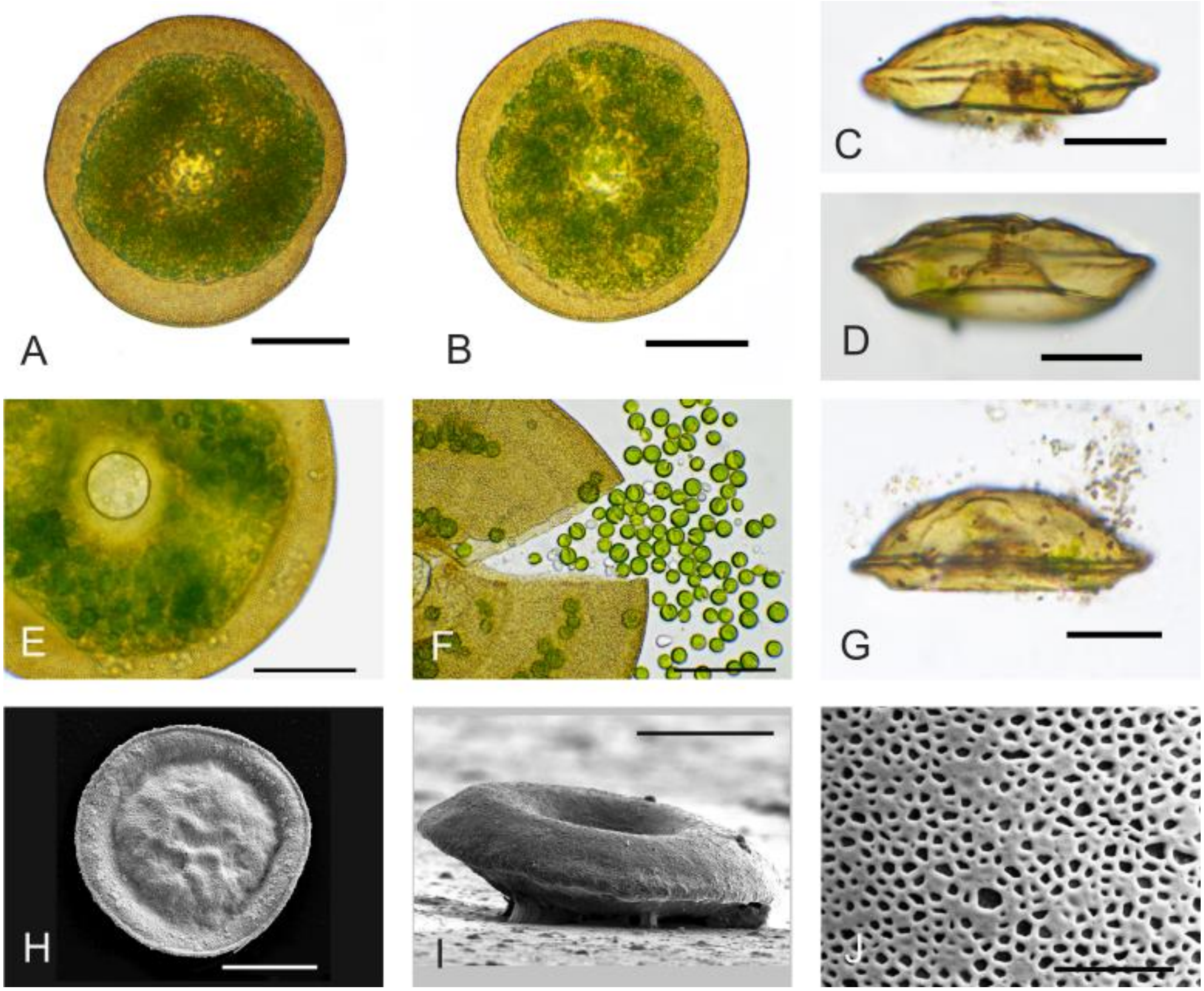


**Supplementary Fig. 5.** A-J. *Arcella artocrea*, from Mer Bleue Bog, Ottawa, Ontario, Canada. A,B. Ventral views; LM, 400×. C,D,G. Profile views; LM, 400×. E. Ventral; LM, 1,000×. F. Ventral of a broken shell, with released endosymbiont algae and granules of glycogen. H. Dorsal, showing a tubular rim around the perimeter of the shell (bourrelet); SEM, 1,000×. I. Profile of an upturned shell, showing the eversion of the basal rim and concavity of the apertural face; SEM, 1,511×. J. Close view of the ventral surface, showing a disorderly arrangement of collapsed areoles; SEM, 10,000×. All scale bars = 50 μm, except J. where scale bar = 5 μm.

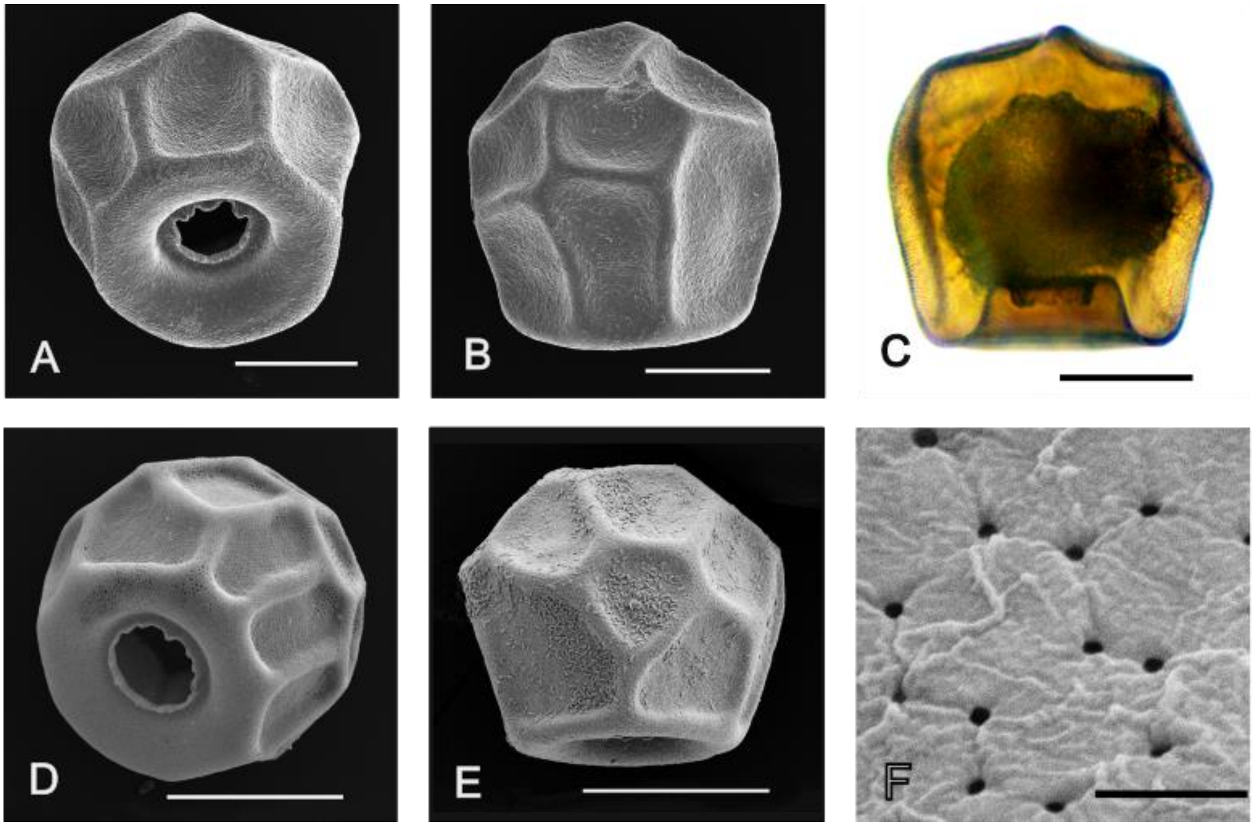


**Supplementary Fig. 6.** A-C. Large morphotype of *Arcella spectabilis*, from Mer Bleue Bog, Ottawa, Ontario, Canada. A. 3/4 apertural view; SEM, 800×. B. Profile; SEM, 800×. C. Profile; LM, 400×. D-F. Small morphotype of *Arcella spectabilis* from Mer Bleue. D. ¾ Apertural view; SEM, 1,318×. E. Profile; SEM, 1,500×. F. Close view of hexagonal building units (areoles). All scale bars = 50 μm, except F. where scale bar = 1 μm.

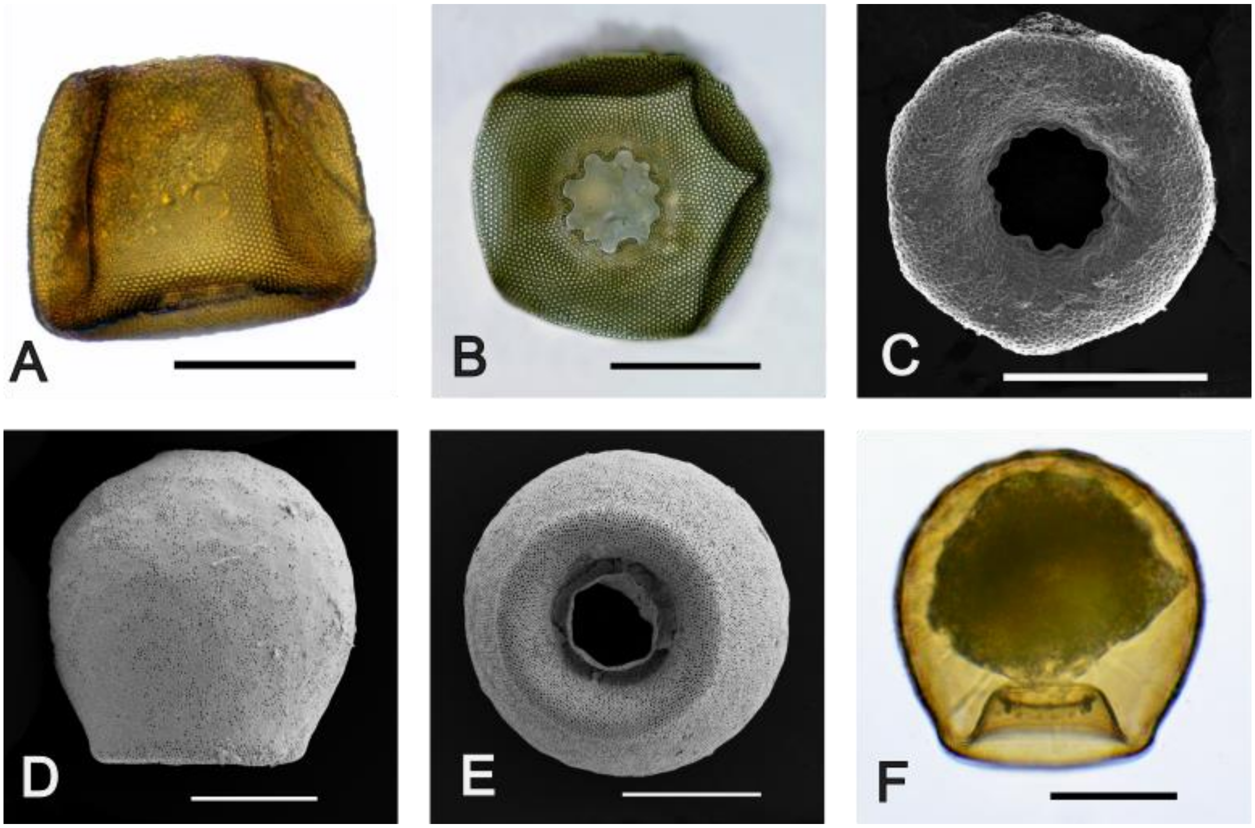


**Supplementary Fig. 7.** A-C. *Arcella prismatica* from Mer Bleue Bog, Ottawa, Ontario, Canada. A. Profile; LM, stacked, 400×. B. Ventral view; LM, 400×. C. Ventral; SEM, 1,500×. D-F. *Arcella mitrata* from Mer Bleue. D. Profile; SEM, 800×. E. Ventral; SEM, 1,000×. F. Profile; LM, 400×. All scale bars = 50 µm.
